# HDAC Inhibitors recapitulate Human Disease-Associated Microglia Signatures *in vitro*

**DOI:** 10.1101/2024.10.11.617544

**Authors:** Verena Haage, John F. Tuddenham, Alex Bautista, Charles C. White, Frankie Garcia, Ronak Patel, Natacha Comandante-Lou, Victoria Marshe, Rajesh Kumar Soni, Peter A. Sims, Vilas Menon, Andrew A. Sproul, Philip L. De Jager

## Abstract

Disease-associated microglia (DAM), initially described in mouse models of neurodegenerative diseases, have been classified into two related states; starting from a TREM2-independent DAM1 state to a TREM2 dependent state termed DAM2, with each state being characterized by the expression of specific marker genes^1^. Recently, single-cell (sc)RNA-Seq studies have reported the existence of DAMs in humans^2–6^; however, whether DAMs play beneficial or detrimental roles in the context of neurodegeneration is still under debate^7,8^. Here, we present a pharmacological approach to mimic human DAM *in vitro* by exposing different human microglia models to selected histone deacetylase (HDAC) inhibitors. We also provide an initial functional characterization of our model system, showing a specific increase of amyloid beta phagocytosis along with a reduction of MCP-1 secretion. Additionally, we report an increase in *MITF* expression, a transcription factor previously described to drive expression towards the DAM phenotype. We further identify *CADM1*, *LIPA* and *SCIN* as DAM- marker genes shared across various proposed DAM signatures and in our model systems. Overall, our strategy for targeted microglial polarization bears great potential to further explore human DAM function and biology.

## Introduction

Initially described in mouse models of neurodegenerative diseases, disease-associated microglia (DAM) have been classified into two related states; cells start from a TREM2 independent DAM1 state to a TREM2 dependent state termed DAM2, with each state being characterized by the expression of specific marker genes^1^. Only recently, single-cell (sc)RNA-Seq studies have reported the existence of DAMs in humans^2–6^. Nevertheless, the question of whether DAM plays a beneficial or detrimental role in the context of neurodegeneration remains a subject of debate^7,8^. To date, only one study has proposed an *in vitro* model system for human DAM based on the exposure of induced pluripotent stem cell-derived microglia-like cells (iMGs) to apoptotic neurons, and the investigators used this model system to assess the phagocytic capacity of these perturbed cells^9^. This model system is a very interesting foray into modeling human DAM but is restricted by the challenge of limited reproducibility by the polarizing agent that is derived from biological material and therefore variable over production batches and tissue sources.

Here, we present an alternative, pharmacological approach to mimic human DAM *in vitro*, prioritizing tool compounds using an *in silico* screening methodology. We show that DAM1-like and DAM2-like signatures exist in single-cell and single-nucleus RNA sequencing datasets derived from human microglia, that we can test prioritized compounds, and that we can reproduce DAM-like transcriptional signatures in human microglia-like cells *in vitro* by exposing different human microglia models to selected histone deacetylase (HDAC) inhibitors. We further provide an initial functional characterization of our model system, elaborating a growing pharmacological toolkit for the microglia community and illustrating our model system’s potential to further explore human DAM biology.

## Results

### Definition of DAM signatures analyzed in this study

We recently generated two datasets derived from single-cell RNA sequencing (scRNAseq) of freshly isolated live primary human microglia derived from 74 donors^6^ and single nucleus RNA sequencing (snRNAseq) profiling of frozen cortex from 424 aging and Alzheimer brains^10^. Both of these independent datasets identified a microglial subtype that is enriched for the DAM2 signature: cluster 11 among the live microglia profiled with scRNAseq^6^ **(Figure 1A**; and microglia 13 in the snRNAseq data^10^ (**Figure 1B**).

**Figure 1.**
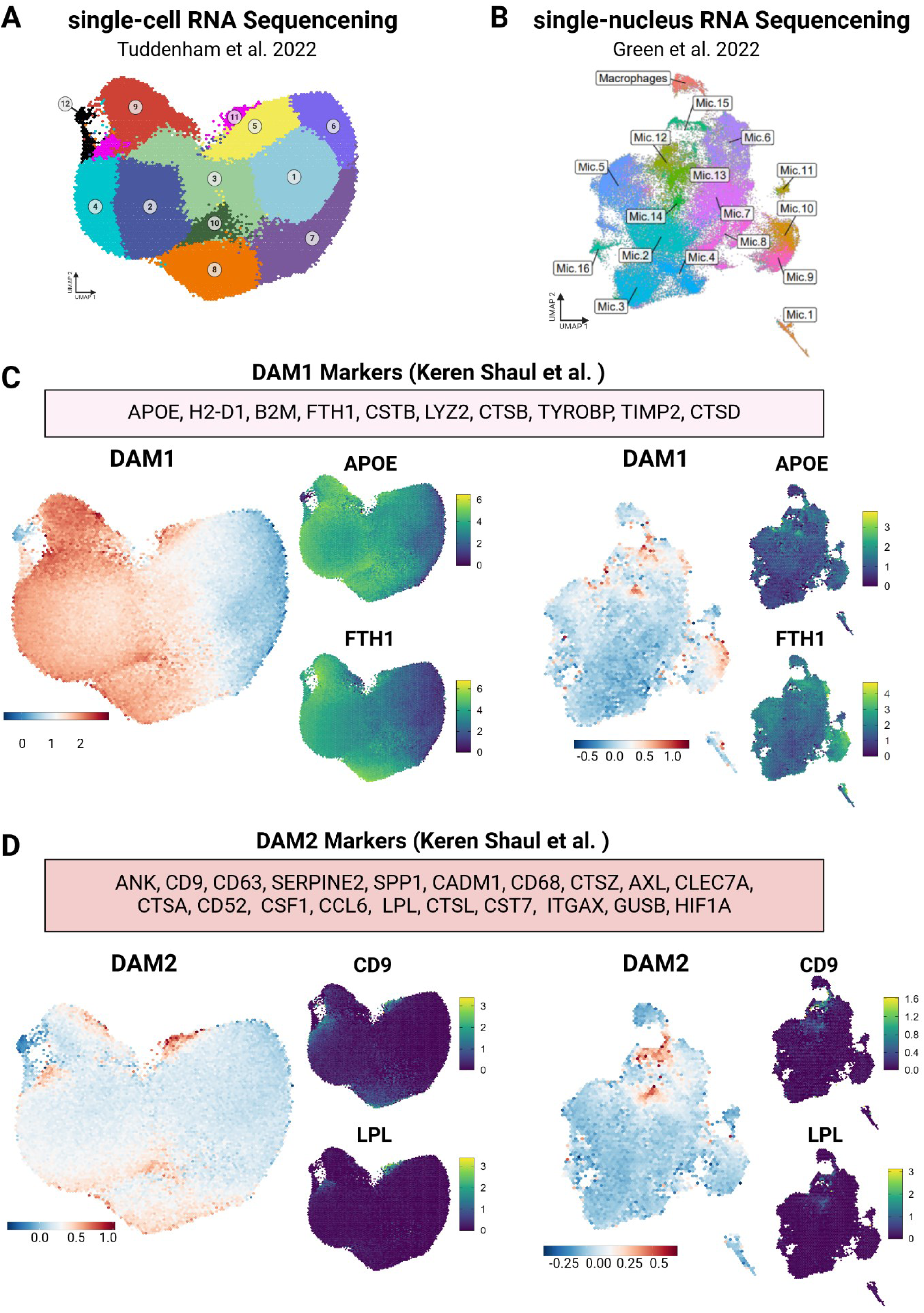
Examining the disease-associated microglial (DAM) signature across human microglia identifies different patterns of capture of DAM-associated genes between single-cell (sc-) and single-nucleus (sn-) RNAseq data. A. UMAP of human single-cell microglial clusters. ^6^. Here, microglia from a single-cell dataset derived from 74 human donors are plotted. 12 microglial clusters were identified. **B. UMAP of human single-nucleus microglial clusters** ^10^. Here, microglia from a single-nucleus dataset derived from 424 human donors are plotted. 16 microglial clusters were identified. **C. DAM1 module expression across the sc- (left) and sn- (right) RNAseq datasets.** Enrichment of the top 10 genes for the DAM1 signature or the top 20 genes for the DAM2 signature from the original publication was calculated on a per-cell basis. Module scores were computed compared to background genes with similar levels of expression. Individual cells are colored by log-fold change of the gene set. Module scores were plotted on hex-binned UMAPs. Individual hexagons are aggregates of 50 cells on average, the plotted score per hexagon is the mean of the score across all cells aggregated within each hexagon. Scores are log-normalized counts, as shown on the color gradient bar. Red/yellow represents the maximal expressed value, while blue/purple represents the lowest expression values. Selected DAM1 (*APOE*, *FTH1*) marker genes were plotted across microglial clusters. **D. DAM2 module expression across the sc- (left) and sn- (right) RNAseq datasets.** Enrichment of the top 10 genes for the DAM1 signature or the top 20 genes for the DAM2 signature from the original publication was calculated on a per-cell basis. Module scores were computed compared to background genes with similar levels of expression. Individual cells are colored by log-fold change of the gene set. Module scores were plotted on hex-binned UMAPs. Individual hexagons are aggregates of 50 cells on average, the plotted score per hexagon is the mean of the score across all cells aggregated within each hexagon. Scores are log-normalized counts, as shown on the color gradient bar. Red/yellow represents the maximal expressed value, while blue/purple represents the lowest expression values. Selected DAM2 (*CD9, LPL*) marker genes were plotted across microglial clusters.

Here, we further evaluate DAM-enriched human microglial subtypes by assessing the expression of a selection of DAM1 (*APOE, H2-D1, B2M, FTH1, CSTB, LYZ2, CTSB, TYROBP, TIMP2, CTSD*) and DAM2 (*ANK, CD9, CD63, SERPINE2, SPP1, CADM1, CD68, CTSZ, AXL, CLEC7A, CTSA, CD52, CSF1, CCL6, LPL, CTSL, CST7, ITGAX, GUSB, HIF1A*) signature genes^1^across the single-cell and single-nucleus human microglial datasets (**Figure 1C-D**). Following gene set enrichment analysis for each set of signature genes (DAM1, DAM2), the log-normalized expression of each set was plotted into the respective UMAP of the human microglial sc- or snRNAseq data. Interestingly, we detected both DAM1 and DAM2 marker gene expression. While the DAM1 signature was more broadly expressed among human microglia, DAM2 expression was restricted to more defined groups of cells, characterized by increased expression of *LPL, LGALS1, CD9* and *GPNMB* (**Figure 1C-D, Figure S1**). These DAM2+ cells include cluster 11 in live microglia, identified as the DAM-enriched cluster in the earlier report^6^.

With regards to the snRNAseq dataset, DAM1 gene expression was more restricted to certain aspects of the distribution of microglia, as was DAM2 (**Figure 1C-D**). Interestingly, as also observed in the scRNAseq data, microglia with DAM2-specific expression were enriched in a region of the distribution of microglia that is high in DAM1 marker expression, suggesting that DAM2 might arise from DAM1, but that not all microglia transition from a DAM1 to a DAM2 state.

At this point, the role of the observed DAM-like signatures needs to be validated. In this large snRNASeq dataset, both the DAM1-enriched and the DAM2-enriched are associated with the amyloid and tau proteinopathies that define AD^10^.

### Identifying compounds that engage the DAM signatures

To establish a model system using human cells, we deployed an *in silico* compound prioritization strategy to identify pharmacological compounds that may either induce or suppress the respective DAM-like signatures from the human datasets with the goal to recapitulate and manipulate those cell subsets *in vitro* and to evaluate their function as we have previously done for other microglial subtypes^11^. In short, we leveraged the Connectivity Map resource (CMAP;^12^), a transcriptomic atlas derived from a range of human cell lines exposed to thousands of pharmacological compounds, to identify molecules that induce or reduce the transcriptomic signature of our DAM-like human microglial subtypes identified from the sc- (Cluster 11^6^, for full signature see **Table S1**) or the snRNAseq datasets (Microglia 13;^10^, **Figure 2A**, for full signature see **Table S1**). Our analysis yielded a specific set of compounds for each of the queried DAM signatures (excerpt of selected compounds in **Figure 2B**, for full list see **Figure S2A and Table S2**). To select candidate compounds, the common upregulated or downregulated genes from the two analyses were merged, and we selected five to six predicted compounds from each comparison based on an absolute tau score >99.5. The focus of this analysis was initially to identify compounds mimicking a more general DAM-like human signature instead of building specific DAM1 and DAM2 *in vitro* models since we did not want to overfit our modeling to the primarily mouse-defined signatures^1^.

**Figure 2.**
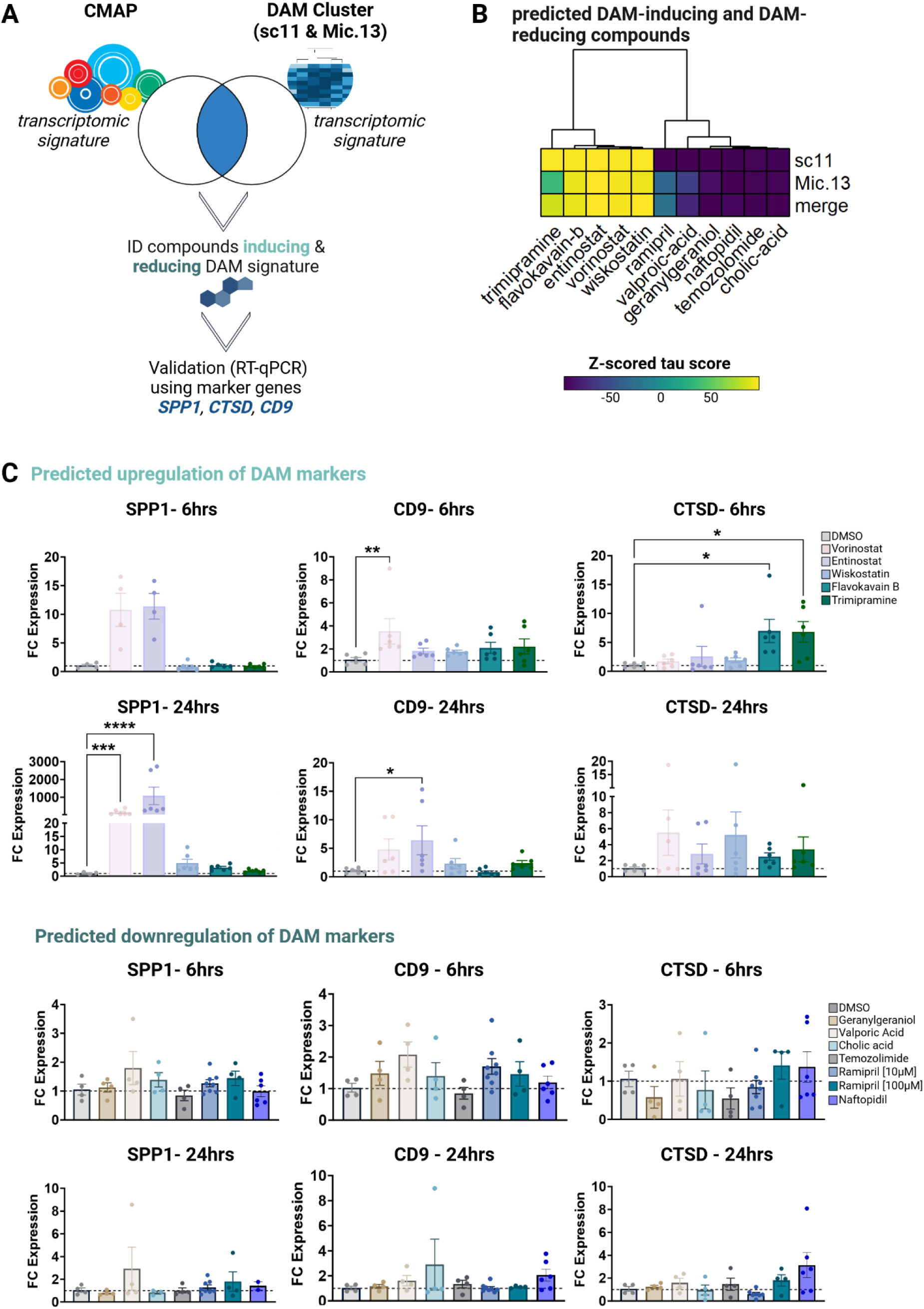
*In silico* compound screen and validation of transcriptomic modulators for the DAM1/DAM2 signature. A. Graph depicting the *in silico* approach to identify compounds mimicking the DAM cluster signatures using the CMAP resource (Connectivity Map resource; ^12^) followed by a validation approach. **B. CMAP predictions** from microglial single-cell RNA- Seq Cluster 11 ^6^, single-nucleus RNA-Seq Microglia 13 cluster ^10^, and the overlapping upregulated gene set. The Connectivity Map (CMAP) was used to identify compounds that are predicted to upregulate or downregulate gene sets associated with either of the DAM-like clusters from single-cell or single-nucleus data. Heatmaps depict the Z-scored tau score for each compound following query analysis. Clustering of tau scores across microglial clusters 11, 13, or the merged signature was performed with absolute linkage. **C. Screening of identified candidate drugs via RT-qPCR in HMC3 microglia-like cells.** HMC3 cells were exposed to selected compounds predicted to upregulate (upper panel; DMSO control: n=6; Vorinostat: n=6; Entinostat: n=6; Wiskostatin: n=6; Flavokavain B: n=6; Trimipramine: n=6;) or downregulate (lower panel; DMSO control: n= 4; Geranylgeraniol: n=4; Valproic acid: n=4 ; Cholic acid: n=4; Temozolomide: n=4; Ramipril: 0.01mM n=8, 0.1mM n=4; Naftopidil: n=6) the DAM signature at 6 hrs. or 24 hrs. Selected DAM marker gene expression (*CD9, SPP1, CTSD*) was assessed via RT-qPCR. CT values were normalized to *HPRT1*. Bars represent fold- change gene expression in relation to DMSO control. For statistical analysis Kruskal Wallis test followed by Dunn’s multiple comparisons test was performed.*p.adj ≤ 0.05; **p.adj ≤ 0.01; ***p.adj ≤ 0.001; ****p.adj ≤ 0.0001.

Our analysis identified a series of intriguing compounds (**Figure 2B**). One of the top families of positive regulators in our screen were histone deacetylase (HDAC) inhibitors, including Entinostat and the FDA-approved Vorinostat, as well as experimental compounds such as Merck60 and APHA-compound-8. Interestingly, except for Vorinostat, which is a pan-HDAC inhibitor, many of these drugs are selective HDAC inhibitors, with HDAC 1/2 being the most common targets for these drugs^13^. As compounds predicted to induce the DAM-like signature we further identified the neural Wiskott-Aldrich syndrome protein (N-WASP) inhibitor Wiskostatin, the tricyclic antidepressant Trimipramine and the hypoxia-inducible factor (HIF) inhibitor Flavokavain B (**Figure 2B**). It is interesting to note that HIF-1α was previously predicted as an upstream regulator of amyloid plaque-associated microglia and has been shown to regulate synaptosomal phagocytosis *in vitro*^14^.

As compounds that downregulate the DAM signature, our *in silico* analysis identified: Geranylgeraniol (an intermediate in the mevalonate pathway), valproic acid (an established treatment for seizures), cholic acid (a naturally occurring bile acid), Ramipril (an angiotensin-converting enzyme (ACE) inhibitor), the prodrug Temozolomide (an alkylating agent currently used in glioblastoma therapy^15^), and Naftopidil (an α1-Adrenoceptor Antagonist) (**Figure 2B**).

### Validation of prioritized compounds in the human HMC3 model system

In order to assess the effect of the selected compounds on the expression of DAM signature genes, the compounds were first titrated on the human microglia cell 3 (HMC3) microglia-like cell line to assess toxicity and to determine doses for each of the compounds that were not toxic to the cells. Specifically, an MTT (3-[4,5-dimethylthiazol-2-yl]-2,5 diphenyl tetrazolium bromide) assay was used and drug concentrations with a comparable absorption to DMSO-treated HMC3 microglia (control condition) were selected for downstream experiments (**Figure S2**). Following the selection of the treatment concentration for each drug, HMC3 microglia were exposed for 6hrs and 24hrs and subsequently, the expression of *CTSD* (DAM1 marker) as well as *SPP1* and *CD9* (DAM2 marker genes) was assessed via RT-qPCR (**Figure 2C**). With regards to the compounds predicted to induce the DAM signature, we identified the two HDAC-inhibitors Vorinostat and Entinostat as our top candidates. We detected significant upregulation of *SPP1* expression, particularly after 24 hrs.: Vorinostat p= 0.0009 and Entinostat p <0.0001. The effect on *CD9* was more modest: 6hrs - Vorinostat p= 0.014 and 24hrs - Entinostat p = 0.038). As both compounds belonged to the class of HDAC inhibitors and showed significant effects, Vorinostat and Entinostat were selected for further validation experiments. HDAC inhibitors have primarily been studied in cancer; however, there is growing interest in their use in the field of neurodegeneration^16^. They have previously been shown to suppress inflammatory responses in microglia^17^.

As it is currently unclear whether DAMs play a beneficial or detrimental role in humans, the identification of compounds with the potential to downregulate the DAM-like signature is also of great interest. However, from our selected candidate compounds none showed a consistent pattern of downregulating the expression of DAM signature genes (**Figure 2C**, lower panel). We therefore proceeded with the DAM-inducing drugs focusing on the development of a human DAM in an *in vitro* model system.

In order to further assess the effect of Vorinostat and Entinostat treatment on the human microglial cell line HMC3, we exposed three independent passages of HMC3 microglia to each of the drugs for 24 hrs. and performed bulk RNA-Seq for each compound. For analysis, we used the DESeq2^18^ package implemented within R (4.4.1) to test for differentially expressed genes between treatment conditions. Subsequently, we assessed the expression of different DAM or DAM-like gene modules, namely DAM1 and DAM2 as defined by Keren-Shaul and colleagues^1^, Cluster 11^6^ and Microglia 13^10^ as defined from recent human datasets (**Table S3A**). To do so, a Wald test was performed to generate a p-value and a resulting Benjamini-Hochberg (FDR) value, where each gene set was analyzed independently from the others (**Figure 3**). After applying this test, any gene with a positive log2 fold-change and an FDR < 0.05 was considered to be statistically significantly upregulated. For a comparison between the different inquired signatures, see **Figure S3A**. For a full list of differentially expressed genes between the different treatment conditions, see **Table S4**.

**Figure 3.**
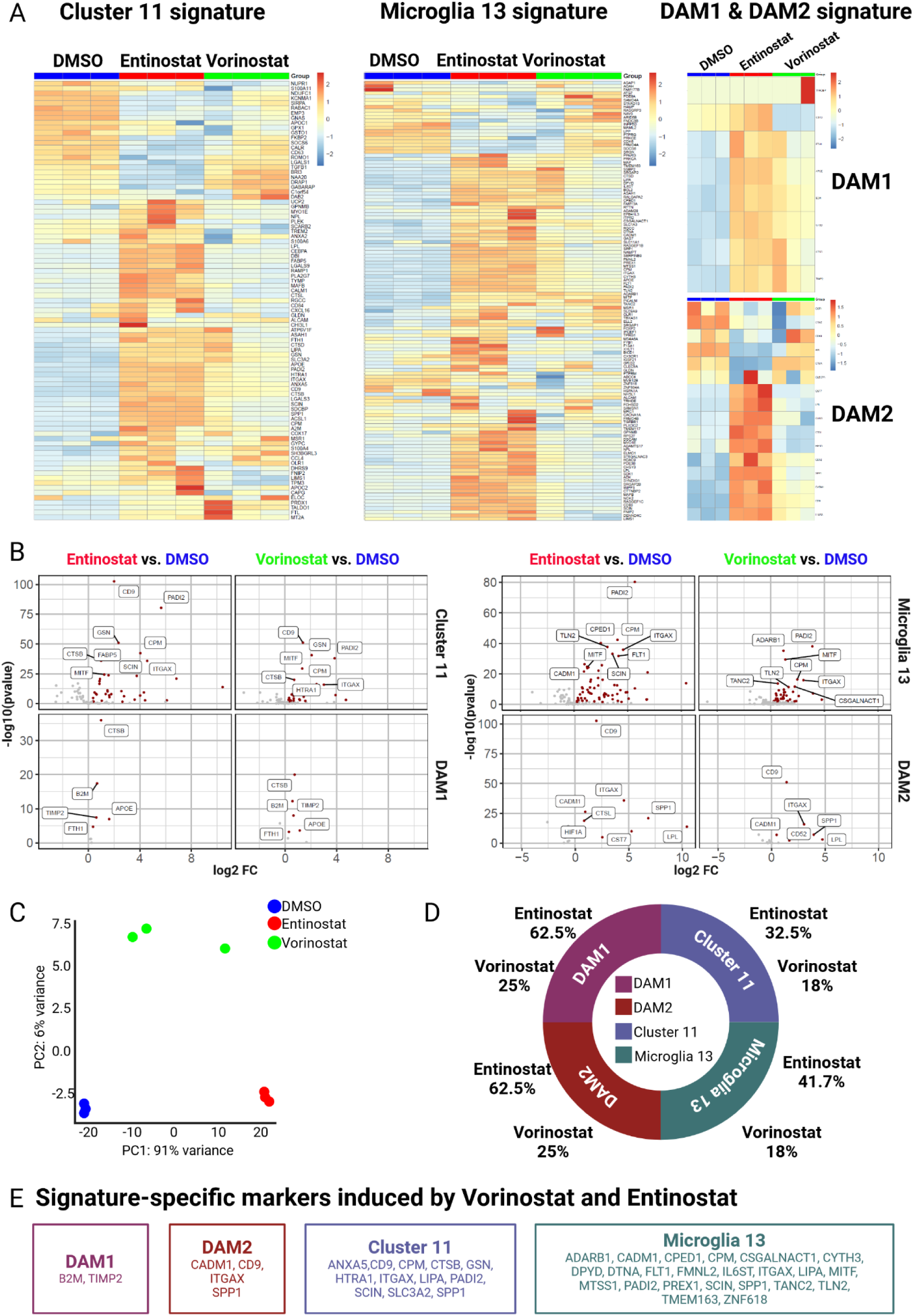
**Bulk RNAseq data from the HMC3 DAM model. A. Heatmaps showing the expression of Cluster 11**^6^ (left), **Microglia 13**^10^ (middle) and **DAM1/DAM2**^1^(right) **marker gene sets** in bulk RNAseq data generated 24hrs following exposure to DMSO (control), Entinostat (red; 10µM) or Vorinostat (green; 1µM). Each column represents a single sample, each row a single gene represented in the respective marker set. Pairwise differential testing between DMSO control and each of the treatment conditions (Entinostat, 10µM; Vorinostat, 1µM) was conducted using a Wald test with the Benjamini-Hochberg correction (FDR alpha < 0.05). The legend represents Z scores, with lower scores indicated in red and higher scores indicated in blue. Data represents n=3 independent experiments for each treatment group with each n for all compounds being performed at the same time. **B. Volcano Plots depicting the distribution of differentially expressed genes from different signatures (DAM1, DAM2**^1^**, Cluster 11**^6^**, Microglia 13**^10^**) for each treatment condition (Entinostat or Vorinostat) in comparison to DMSO control.** HMC3 microglia were treated for 24hrs with DMSO as control, Entinostat (10µM) or Vorinostat (1µM) followed by bulk RNA-Seq. Volcano plots depict all genes present in each marker set (DAM1: 10 genes; DAM2: 20 genes; Cluster 11: 89 genes, Microglia 13: 127 genes) plotted based on log2FC (fold change expression) and -log10(p value) with the ones significantly upregulated marked in red and labelled with the gene name. Plots are organized from Cluster 11 (top left), to DAM1 (bottom left), to DAM2 (top right) to Microglia 13 (bottom right). **C. PCA plot of bulk RNAseq results from HMC3 microglia treated with DMSO, Vorinostat or Entinostat**. Principal component analysis (PCA) was calculated on log-normalized bulk RNA-Seq data derived from compound-treated HMC3 microglia following 24hrs of exposure to DMSO (control; blue), Entinostat (10µM; red) or Vorinostat (1µM; green). Data represents n=3 independent experiments for each of the treatment group with each n for all compounds being performed at the same time. **D. Pie chart depicting the number of significantly upregulated genes by Entinostat or Vorinostat for each of the queried marker signatures DAM1, DAM2, Cluster 11**^6^**, Microglia 13**^10^. The number of significantly upregulated genes across all three replicates for each treatment group (Entinostat or Vorinostat) in comparison to DMSO control was identified and converted to a percentage of marker genes upregulated/ marker set. Data for DAM1 are depicted in purple, for DAM2 depicted in red, for Cluster 11 depicted in violet and for Microglia 13 depicted in teal. **E. Signature-specific markers induced by Vorinostat and Entinostat.** Markers significantly induced by Vorinostat and Entinostat for each signatured are depicted for DAM1 (purple), DAM2 (red), Cluster 11 (violet), Microglia 13 (teal).

Following Principal Component analysis (PCA), Entinostat-treated cell samples clustered closely together and were distinct from DMSO-treated cells, while Vorinostat-treated samples clustered between the two other conditions (**Figure 3C**). When assessing the Cluster 11 signature, interestingly we noticed that a portion of these markers was expressed at baseline by DMSO control cells (**Figure 3A**). While Vorinostat significantly induced the expression of 16/89 (18%) Cluster 11 marker genes, Entinostat exposure engaged a broader set of genes, significantly inducing 29/89 (33%) markers that are not expressed under the control condition (DMSO). These data clearly suggest that the DAM signatures are composed of at least two sets of genes whose transcriptional regulation is somewhat distinct: these two sets of genes may be co-regulated in the contexts where they were derived, but our precise molecular perturbation reveals a difference in regulation of the two gene sets, one of which seems to be engaged at baseline by the culture system. Overall, *CD9*, *PADI2*, *GSN* and *CTSB* (**Figure 3B**) were genes strongly induced by both compounds. *CD9* is a key DAM marker gene, while *PADI2* has been associated with neurodegeneration in microglia^19^. *CTSB* has been reported as a potential major driver in brain aging^20^.

Interestingly, when assessing the expression of the Microglia 13 marker signature^10^, we observed a minor subset of these signature genes being expressed at baseline conditions (DMSO controls), and they were downregulated upon Entinostat treatment and slightly reduced upon Vorinostat exposure (**Figure 3A**). However, the majority of the Microglia 13 gene signature is engaged by our HDAC inhibitors: Entinostat potently and significantly induced 53/127 of Microglia 13 genes, which is 42% of the signature. Similar to the results from Cluster 11, Vorinostat significantly induced a lower percentage of genes belonging to the Microglia 13 signature (23/127 genes; 18% of the signature). While Vorinostat and Entinostat both induced *PADI2 (*Vorinostat - padj =3.69E^-36^; Entinostat - padj = 2.97E^-78^) most significantly, they also both induced *MITF (*Vorinostat – padj = 9.01E^-28^; Entinostat – padj = 1.19E^-23^), a transcription factor recently shown to be an important driver of the DAM signature and a highly phagocytic phenotype in human iPSC-derived microglia-like cells^9^ (**Figure 3B**).

When assessing the expression of the originally defined murine signatures of DAM1 and DAM2^1^, we also observed that a small fraction of those genes are expressed by DMSO-treated control cells, and we see a strong and significant induction of both signatures by Entinostat (E): about 63% of each signature is engaged (DAM1: 5/8 genes; DAM2:10/16 genes). On the other hand, Vorinostat (V) significantly induced about 38% of the DAM1 (3/8 genes) and 25% of the DAM2 signature genes (4/16 genes). Both compounds induced *B2M* and *TIMP2* as DAM1 markers and jointly induced the DAM2 markers *SPP1*, *ITGAX*, *CD9*, *CADM1* (**Figure 3E**; **Table 1**). **Figure 3D** depicts an overview of the number of markers from each signature induced by each of the compounds (Vorinostat, Entinostat). **Figure 3E** provides an overview of signature-specific markers induced by both compounds, Vorinostat and Entinostat.

**Table 1.** Summary of marker genes jointly induced by Vorinostat- and Entinostat- treatment in HMC3 microglia.

| Gene | Signature | Vorinostat treatment<br>[p.adj; log2FC] | Entinostat treatment (p.adj)<br>[p.adj; log2FC] |
| --- | --- | --- | --- |
| *PADI2* | Cluster 11 & Mic13 | 3.69E<sup>-36</sup> [3.87] | 2.97E<sup>-78</sup> [5.66] |
| *SPP1* | DAM2 | 8.65E<sup>-07</sup> [3.94] | 2.39E<sup>-20</sup> [6.85] |
| *ITGAX* | DAM2 | 7.90E<sup>-15</sup> [3.04] | 1.12E<sup>-34</sup> [4.57] |
| *CD9* | DAM2 | 5.16E<sup>-49</sup> [1.41] | 3.47E<sup>-100</sup> [2.00] |
| *MITF* | Mic13 | 9.01E<sup>-28</sup> [1.35] | 1.19E<sup>-23</sup> [1.23] |
| *B2M* | DAM1 | 2.16E<sup>-11</sup> [0.56] | 5.77E<sup>-17</sup> [0.68] |
| *TIMP2* | DAM1 | 1.58E<sup>-07</sup> [0.64] | 1.57E<sup>-07</sup> [0.62] |
| *CADM1* | DAM2 | 1.16E<sup>-06</sup> [0.48] | 1.62E<sup>-25</sup> [0.96] |

Overall, our two prioritized HDAC inhibitors engage overlapping aspects of the DAM signatures; however, these signatures appear to be complex, consisting of at least 2 sets of genes with distinct transcriptional programs, one of which is upregulated in the DMSO control condition. The second, larger gene set includes the key marker genes and is engaged by these compounds. This is not surprising as microglia and microglia-like cells are highly reactive and are unlikely to be in a homeostatic state in culture^21^. Nonetheless, the HDAC inhibitors clearly engage an important component of the DAM signatures, and these signatures need to be refined to guide future study designs.

### Evaluation of Vorinostat in the iMG model system

We additionally tested one of the DAM-inducing compounds - Vorinostat- on iPSC-derived microglia (iMG) on Day 28-29 of the iMG differentiation protocol. Following 24 hours of exposure to the compound, iMG were harvested and subjected to bulk RNA-Seq profiling followed by an analysis assessing the expression of five different marker sets (**Figure 4**). In addition to the DAM1, DAM2, Cluster 11 and Microglia 13 signatures tested above in control- (DMSO) vs. Vorinostat-treated cells^1,6,10^, we also assessed the expression of signature genes recently derived from an iMG-derived human DAM model based on exposure to a preparation of apoptotic neurons^9^ (**Figure S4A; Table S3B**). The authors of that report identified two cell clusters related to the DAM subtype, which they termed Cluster 2 and 8. As a result, we refer to this signature from now on as iMG Cluster 2+8. In our analysis, we subsequently put a major focus on the signatures defined from human data, namely Cluster 11^6^, Microglia 13^10^ and iMG Cluster 2+8^9^. Data from samples derived from one iMG batch (n= 5 replicates) and the treatment conditions (DMSO, Vorinostat) are presented in **Figure S4B**. **Figure 4A** depicts all significantly induced genes from each of the queried signatures in red, while a selection of the nine most upregulated genes for each signature is highlighted through labelling. As in the HMC3 model system, we observed some baseline expression of marker genes across all signatures in DMSO-treated control iMG (**Figure 4A-B**). Vorinostat significantly induced 12% of the Cluster 11 and Microglia 13 signature as well as 15% of the iMG Cluster 2+8 signature across all replicates. With regards to Cluster 11^6^ genes, the most significantly induced genes were *DHRS9*, *RABAC1 and NPL*. With regards to the Microglia 13 marker signature^10^, *ARGHAP48*, *PTPRG* and *SCIN* were the most significantly induced markers, and, for the Cluster 2+8 signature^9^, *CYSTM1, ABCA1* and *SLC38A6* were most significantly upregulated (**Figure 4A**; **Table 2**). When assessing the classical DAM1 and DAM2 markers, Vorinostat did not induce any DAM1 and about 20% of DAM2 marker genes (**Figure S4C**). We then generated a second batch of iMGs and replicated most of our findings (**Figure S4D-F**) following exposure to Vorinostat. For a full list of differentially expressed genes between the different treatment conditions, see **Table S4**.

**Figure 4.**
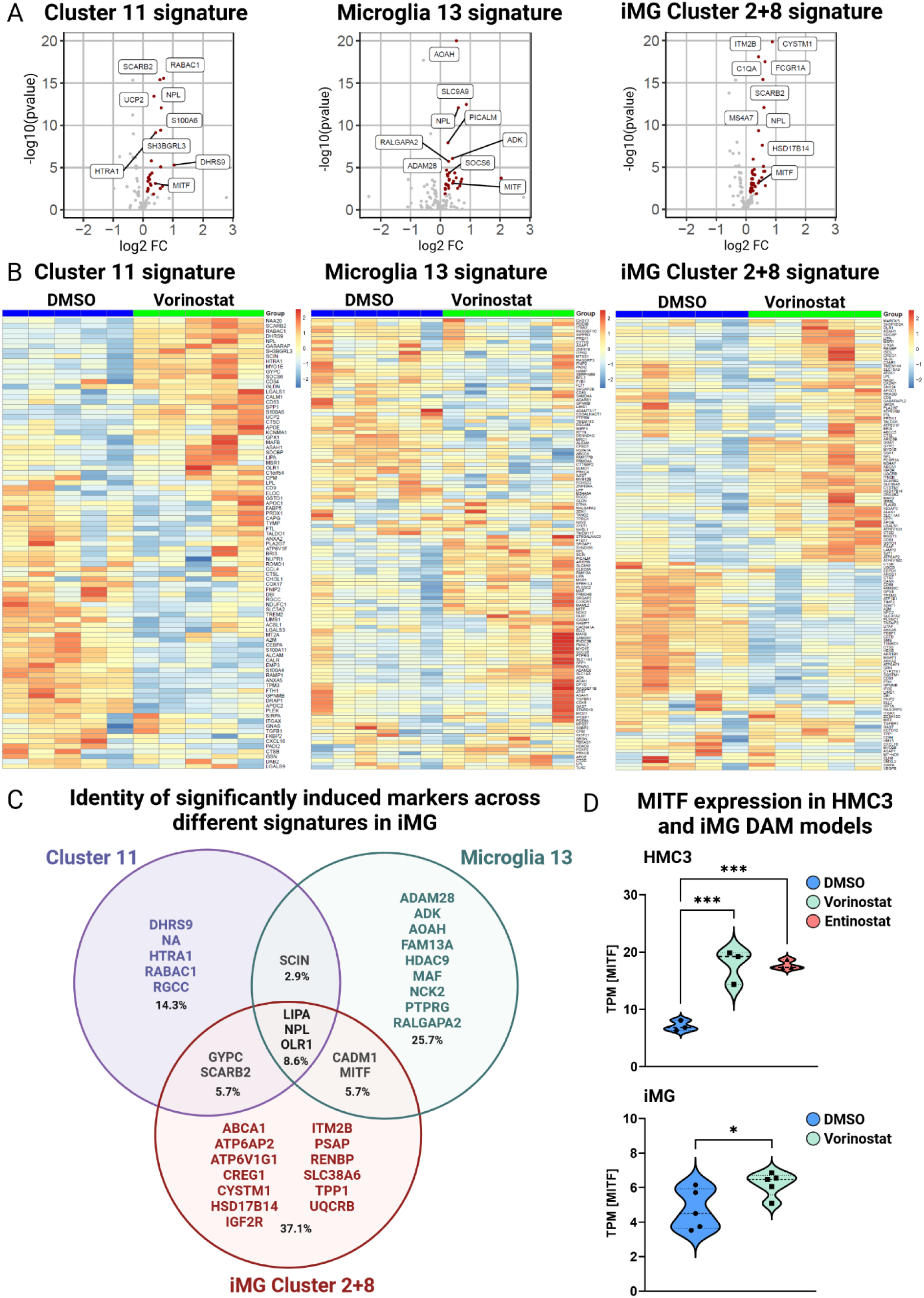
**Bulk RNA-Seq of the human iPSC-derived microglia (iMG) DAM model. A. Volcano Plots depicting the distribution of differentially expressed genes from different signatures (Cluster 11**^6^**, Microglia 13^10^, iMG Cluster 2+8**^9^**) for Vorinostat treatment in comparison to DMSO control.** iPSC-derived microglia at Day 28-29 of differentiation were treated for 24hrs with DMSO as control or Vorinostat (0.1µM) followed by bulk RNAseq. Volcano plots depict all genes present in each marker set (Cluster 11: 89 genes, Microglia 13: 127 genes, iMG Cluster 2+8: 134 genes) plotted based on log2FC (fold change expression) and -log10(p value) with the ones significantly upregulated marked in red and of the most significantly changed genes, a selection of nine genes was labeled with the gene name. Plots are organized from Cluster 11 (left), to Microglia 13 (middle), to iMG Cluster 2+8 (right). **B. Heatmaps showing the expression of Cluster 11**^6^ (left), **Microglia 13**^10^ (middle) and iMG Cluster 2+8^9^ (right) **marker sets** in bulk RNAseq data generated 24hrs following compound treatment with DMSO (control) or Vorinostat (green; 0.1µM). Each column represents a single sample, each row a single gene represented in the respective marker set. Pairwise differential testing between DMSO control and each of the treatment conditions (Entinostat, 10µM; Vorinostat, 1µM) was conducted using a Wald test with the Benjamini-Hochberg correction (FDR alpha < 0.05). The legend represents Z scores, with lower scores indicated in red and higher scores indicated in blue. Data represents n=5 independent experiments per treatment group from one batch of iPSC-derived human microglia. For data replication in a second batch see Supple. Fig. 4. **C. Venn diagram depicting significantly induced markers across the signatures for Cluster 11**^6^**, Microglia 13**^10^ **and iMG Cluster 2+8**^9^ **in Vorinostat-treated iMGs.** Each circle shows significantly induced markers from each marker set - Cluster 11 (violet), Microglia 13 (green), Dolan et al. (red). Overlays of circles depict induced marker genes shared across different combinations of marker sets. Percentage indicates ratio of each marker set in relation to the total number of significantly induced markers across all three signatures. **D. MITF expression in HMC3 and iMG DAM models.** Violin plots depict the expression of the transcription factor MITF in transcripts per million (TPM) across treatment conditions in HMC3 microglia (top; DMSO (blue), Vorinostat (1µM; green), Entinostat (10µM; red); n=3/group) and iMG (bottom; DMSO (blue), Vorinostat (0.1µM; green); n=6 per group, one iMG batch; for data replication see Suppl. Fig.4). For statistical analysis of HMC3 data, one-ay ANOVA followed by Dunnett’s multiple comparisons test was performed. For iMG data, unpaired t-test was performed. Each dot represents a replicate, central interrupted line represents the median and fine dotted lines represent the interquartile range. *p.adj ≤ 0.05, **p.adj ≤ 0.01, ***p.adj ≤ 0.001 test

**Table 2.** Summary of marker genes induced by Vorinostat treatment in iMG.

| Gene | Signature | Vorinostat treatment<br>[p.adj; log2FC] |
| --- | --- | --- |
| *DHRS9* | Cluster 11 | 5.68E<sup>-05</sup> [1.04] |
| *RABAC1* | Cluster 11 | 1.77E<sup>-14</sup> [0.69] |
| *ARGHAP48* | Mic13 | 1.68E<sup>-03</sup> [0.71] |
| *PTPRG* | Mic13 | 3.14E<sup>-03</sup> [0.69] |
| *SCIN* | Mic13 + Cluster 11 | 1.40E<sup>-02</sup> [0.58] |
| *CYSTM1* | iMG Cluster 2+8 | 1.55E<sup>-18</sup> [0.83] |
| *ABCA1* | iMG Cluster 2+8 | 2.83E<sup>-04</sup> [0.63] |
| *SLC38A6* | iMG Cluster 2+8 | 2.96E<sup>-04</sup> [0.59] |
| *LIPA* | Cluster 11/Mic3/ iMG Cluster 2+8 | 5.92E<sup>-04</sup> [0.23] |
| *NPL* | Cluster 11/Mic3/ iMG Cluster 2+8 | 3.37E<sup>-11</sup> [0.60] |
| *OLR1* | Cluster 11/Mic3/ iMG Cluster 2+8 | 1.41E<sup>-02</sup> [0.16] |

Commonly induced across all three signatures were the markers *LIPA*, *NPL* and *OLR1* (**Figure 4C**, **Table 2**). While *CADM1* and *MITF* are induced markers shared between Microglia 13 and the iMG Cluster 2+8 signature, *GYPC* and *SCARB2* are induced markers shared between the Cluster 11 and iMG Cluster 2+8 signature and *SCIN* was the only induced marker shared between the Cluster 11 and Microglia 13 signature (**Figure 4C**). For an overview of the signature genes for each signature induced by Vorinostat across HMC3 microglia and iMG, see **Figure S4G**. As *LIPA*, *CADMI1*, *MITF* and *SCIN* were induced across both, HMC3 and iMG model systems and are derived from DAM signatures defined from human datasets, we conclude that our HDAC-inhibitor-based model constitutes a robust and valid system to study aspects of human DAM (**Table 3**). For a full list of differentially expressed genes between the different treatment conditions, see **Table S4**.

**Table 3.** Summary of marker genes induced by both compounds in HMC3 and iMG (only Vorinostat).

| Gene | Signature | Entinostat HMC3<br>[p.adj; log2FC] | Vorinostat HMC3<br>[p.adj; log2FC] | Vorinostat iMG<br>[p.adj; log2FC] |
| --- | --- | --- | --- | --- |
| *LIPA* | Cluster 11, Mic13, iMG 2+8 | 5.68E<sup>-11</sup> [1.48] | 2.25E<sup>-07</sup> [1.22] | 5.92E<sup>-04</sup> [0.23] |
| *CADMI1* | Mic13, iMG 2+8 | 1.62E<sup>-25</sup> [0.96] | 1.16E<sup>-06</sup> [0.48] | 5.72E<sup>-03</sup> [0.18] |
| *MITF* | Mic13, iMG 2+8 | 1.19E<sup>-23</sup> [1.23] | 9.01E<sup>-28</sup> [1.35] | 4.52E<sup>-03</sup> [0.41] |
| *SCIN* | Cluster 11, Mic13 | 4.79E<sup>-32</sup> [3.57] | 1.56E<sup>-05</sup> [1.47] | 1.40E<sup>-02</sup> [0.58] |

**Table 4.** Overview of identified DAM-mimicking compounds.

| Compound | Vendor | Cat# | Cluster signature | Concentration used for *in vitro* experiments |
| --- | --- | --- | --- | --- |
| Vorinostat | Ambeed | A234507 | 11 (UP) | 1μM |
| Entinostat | Ambeed | A122285 | 11 (UP) | 10μM |
| Wiskostatin | Cayman chemical | 15047 | 11 (UP) | 1μM |
| Naftopidil | APExBIO | 57149-07-2 | 11 (UP) | 10μM |
| Flavokavain B | ChromaDex | ASB-00006058-005 | 11 (UP) | 10μM |
| Trimipramine | Cayman chemical | 15921 | 11 (UP) | 10μM |
| Amiodarone hydrochloride | Sigma | A8423 | 11 (UP) | 10μM |
| Geranylgeraniol | Sigma | 24034-73-9 | 11 (DOWN) | 10μM |
| Valporic acid | Sigma | PHR1061 | 11 (DOWN) | 0.9μg/ml |
| Ramipril | APExBIO | B2208 | 11 (DOWN) | 10μM |
| Cholic acid | Cayman chemical | 20250 | 11 (DOWN) | 10μM |
| Temozolomide | APExBIO | B1399 | 11 (DOWN) | 100μM |

### *MITF* expression in the DAM model systems

We additionally specifically focused on *MITF* expression in our DAM models using HMC3 microglia and iMGs, as *MITF* has been recently suggested to drive a DAM signature and a phagocytic phenotype^9^ (**Figure 4D**). Vorinostat and Entinostat induced a highly significant increase in *MITF* expression in HMC3 microglia (One-way ANOVA followed by Dunnett’s multiple comparisons test; Vorinostat: p= 0.0008; Entinostat: p= 0.0008). Moreover, the Vorinostat-induced increase in *MITF* expression was also observed in our iMG-DAM model (Unpaired t-test, Vorinostat - p=0.0406). Thus, the effect appears to be preserved across the different DAM model systems. These data strengthen the validity of our model system with regards to prior reports of the role of *MITF* in driving a disease-associated microglia like signature and a highly phagocytic phenotype^9^ and position our compound-based approach as a highly reproducible system to study human DAM and refine the role of the signature’s component genes.

### Functional characterization of the HDAC-inhibitor induced *in vitro* DAM model

As one of the main goals of establishing an *in vitro* model for human DAM is the study of their functional properties, we deployed functional assays next. To assess the phagocytic phenotypes of our compound-driven models of human DAM, HMC3 microglia-like cells were pretreated with Vorinostat, Entinostat or DMSO (control) for 24 hrs., followed by exposure to three distinct substrates: pHrodo Dextran to monitor macropinocytosis, fluorescently labeled Aβ to assess a phagocytic phenotype relevant to amyloid proteinopathy as well as pHrodo-labeled Escherichia coli (E. coli) to assess a phagocytosis associated with acute neuroinflammation. Flow-cytometry was used as a readout (**Figure 5A**). As a control, we pretreated the cells with Cytochalasin D (**Figure S5A**). When assessing macropinocytosis through pHrodo Dextran uptake, we observed an upregulation in both Vorinostat- and Entinostat-treated HMC3 cells, with Vorinostat showing a slightly higher increase (**Figure 5B**). When assessing the uptake of Aβ as model of phagocytosis in a neurodegenerative context, both compounds also showed an increase in Aβ uptake in comparison to DMSO control with Vorinostat (p= ≤ 0.0001) inducing again a more pronounced effect than Entinostat (p= 0.0274)(**Figure 5C**). Interestingly, Aβ can be taken up via macropinocytosis and phagocytosis^22^. Thus, our two HDAC inhibitors revealed a specialization towards an increased uptake of soluble substrates, as E.coli phagocytosis was significantly decreased in Vorinostat-treated cells. Entinostat-treated cells did not show a significant decrease in E. coli uptake, although there was a trend in that direction (**Figure 5D)**.

**Figure 5.**
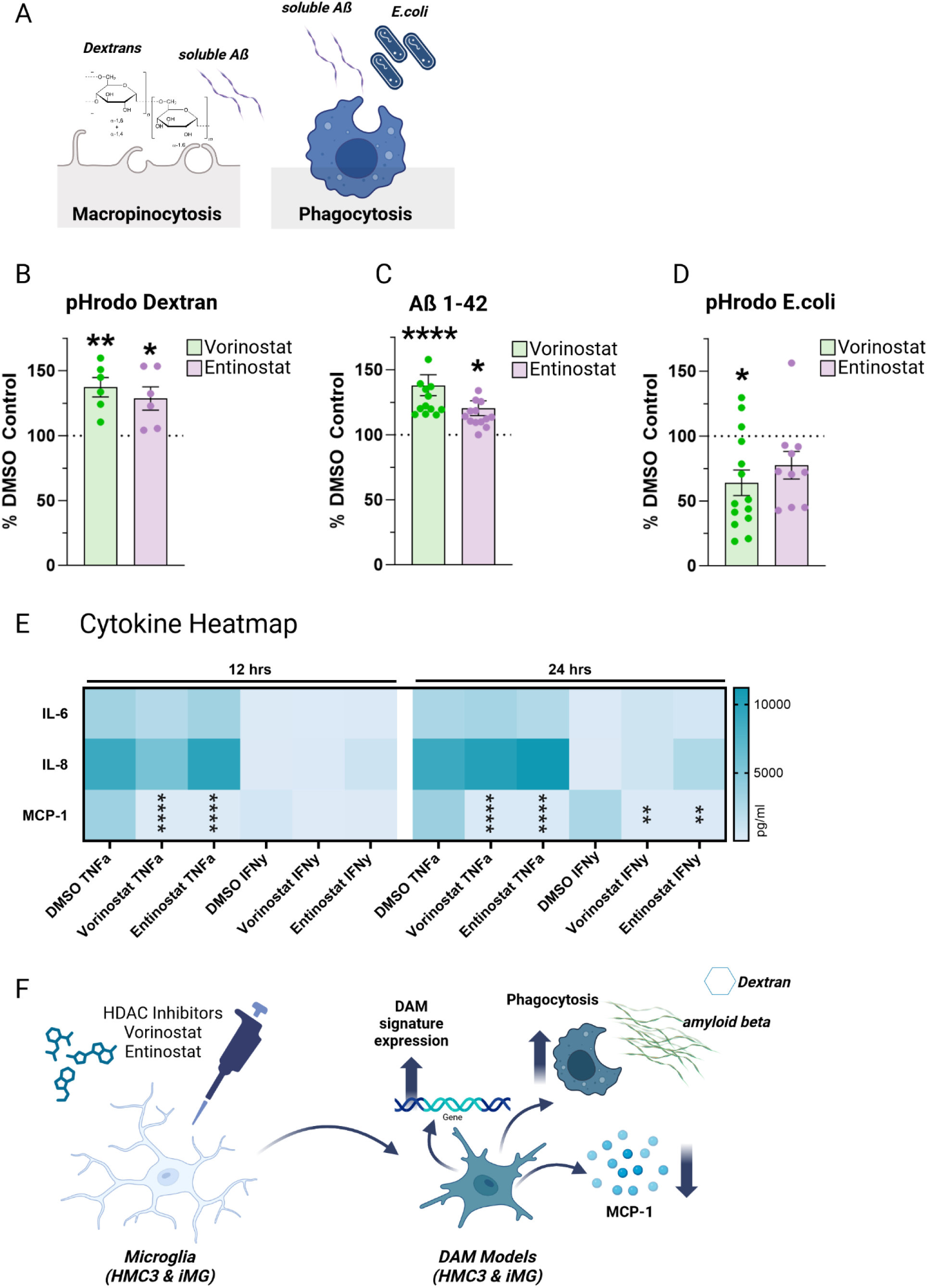
Compound-treated HMC3 microglia exhibit substrate-specific endocytic and phagocytic phenotypes as well as differences in secretion of pro-inflammatory cytokines. A. Graph depicting different assays assessing macropinocytosis (pHrodo Dextran, soluble Aß1-42) and phagocytosis (Aß1-42, E.coli). B. Vorinostat and Entinostat upregulate pHrodoDextran phagocytosis. HCM3 microglia-like cells were pretreated each compound or DMSO as control for 24hrs. Subsequently, they were exposed to pHrodo-labeled Dextran for 1hr, and subsequently the uptake of pHrodo-labeled Dextran was assessed using flow cytometry. Individual experiments are depicted as individual dots in the bar graphs depicting mean ± SEM (Vorinostat – green; Entinostat– purple). Phagocytosis was normalized to percent DMSO (% DMSO) control and for statistical analysis, log-fold change values in comparison to DMSO-treated control samples were analyzed using one-way ANOVA followed by Dunnett’s multiple comparison test. *p.adj ≤ 0.05; **p.adj ≤ 0.01. **C. Vorinostat and Entinostat induce Aß1-42 phagocytosis in HMC3 microglia-like cells.** HCM3 microglia were pretreated with each compound or DMSO as control for 24hrs, subsequently exposed to AlexaFluor 647-labeled Aß monomers for 1hr and subsequently the uptake of AlexaFluor 647-labeled Aß monomers was assessed using flow cytometry Individual experiments are depicted as individual dots in the bar graphs depicting mean ± SEM (Vorinostat - green; Entinostat - purple). Phagocytosis was normalized to % DMSO control and for statistical analysis, log-fold change values in comparison to DMSO-treated control samples were analyzed using one-way ANOVA followed by Dunnett’s multiple comparison test. *p.adj ≤ 0.05; ** p.adj ≤ 0.01. **D. Vorinostat reduces phagocytosis of pHrodo-labeled E.coli.** HCM3 microglia-like cells were pretreated with each compound or DMSO as control for 24hrs, subsequently exposed to pHrodo-labeled E.coli particles for 1hr and subsequently the uptake of pHrodo-labeled E.coli particles was assessed using flow cytometry. Individual experiments are depicted as individual dots in the bar graphs depicting mean ± SEM (Vorinostat - green; Entinostat - purple). Phagocytosis was normalized to % DMSO control and for statistical analysis, log-fold change values in comparison to DMSO-treated control samples were analyzed using one-way ANOVA followed by Dunnett’s multiple comparison test. *p.adj ≤ 0.05; **p.adj ≤ 0.01. **E. Vorinostat and Entinostat, decrease the secretion of pro-inflammatory cytokine MCP-1.** HMC3 microglia-like cells were pre-treated with Vorinostat or Entinostat for 24hrs, and subsequently stimulated with either TNF-α (0.3 µg/mL), IFN-y (0.3 µg/mL) or H2O as control for 12 or 24 hrs. Supernatant was collected and pro-inflammatory cytokine secretion assessed using a human pro-inflammatory cytokine discovery assay. Heatmap depicts measured amount of cytokines IL-6, IL-8, MCP-1 (mean ± SEM) in pg/ml for DMSO control-treated samples (white, light gray, gray) or compound-treated samples (light green, green dark green/light purple, purple, dark purple). For statistical analysis, one-way ANOVA followed by Tukey’s multiple comparisons test with a single pooled variance was performed. *p.adj ≤ 0.05; **p.adj ≤ 0.01; ***p.adj ≤ 0.001; ****p.adj ≤ 0.0001. **F. Summary graph** depicting overall results from this study.

As communication, amplification and orchestration of the immune response is mediated by cytokines, we also assayed the response of our model system to stimulation with one of two pro-inflammatory cytokines that play an important in brain and systemic inflammation: cytokine secretion was evaluated following TNF-α or IFN-γ stimulation after 12 and 24 hours (**Figure 5E**). A panel of 15 cytokines was used as the outcome measure, the secretion of 14 of the cytokines – illustrated by IL-6 and IL-8 – remained unaffected, but both Vorinostat- and Entinostat-treatment significantly (p<0.0001) reduced monocyte chemoattractant protein-1 (MCP-1)(also known as chemokine (C-C motif) ligand 2, CCL2) secretion. This effect was more pronounced with TNF-α stimulation, as the effect persisted over 24 hrs. Thus, our two HDAC inhibitors appear to have a relatively targeted effect on an inflammatory response, one that involves the recruitment of myeloid cells and other leukocytes.

## Discussion

This report presents an approach to study microglia-like cells that express key elements of the human DAM signatures in a reproducible fashion. This is essential to enable the functional characterization of this subtype of microglia *in vitro* in a standardized manner over time and across laboratories; functional characterization of primary microglia is impractical with current technologies given the difficulty of accessing a reasonable number of these cells from the human brain. Our *in silico* analyses identified HDAC inhibitors as a class of molecules with the potential to recapitulate the signatures seen in this subtype of microglia that has been associated with neurodegenerative disease in humans and mice^1,2,4–6^. We prioritized two of these molecules, Vorinostat and Entinostat, and validated our prediction, showing (1) recapitulation of key aspects of the signatures derived from primary human microglia, (2) induction of *MITF*, recently proposed as a regulator of this signature in a study using a different, less reproducible polarization strategy^9^, (3) substrate-specific alterations of uptake consistent with the proposed enhanced phagocytic capacity of DAM-enriched cells^1,9^, and (4) a targeted reduction of MCP-1 secretion following TNF-α or IFN-γ stimulation.

The two HDAC inhibitors - Vorinostat and Entinostat - yield a reproducible model system, that captures aspects of the various murine (DAM1, DAM2) and human (Cluster 11) signatures, but its transcriptional effect resembles most closely the recently identified human Microglia 13 signature that is proposed to mark a microglial subtype contributing to the accumulation of AD pathology^10^. Further, we describe sharing of marker genes and functional changes (increased uptake) with a prior effort to model these signatures *in vitro* using apoptotic neurons as a polarizing agent in iMG towards a DAM-like phenotype^9^. We therefore have addressed the challenge of reproducibility that is intrinsic to the use of cell-preparation derived reagents. In addition, Vorinostat and Entinostat are well-characterized tool compounds that can serve as reference molecular structures for further optimization of desired compound characteristics. Moreover, Vorinostat is approved by the Federal Drug Administration for the treatment of cutaneous T cell lymphoma^23^, facilitating translation to human study participants.

Our study has certain limitations; first, given the difficulty of accessing primary human microglia, we used cellular model systems in our experiments. The usage of two distinct human microglia-like models - the HMC3 cell line and iMG - mitigates this limitation: the two model systems are complementary and provide consistent results. Further, since the original DAM signature is derived from mice and there is no single, generally accepted human DAM signature, we elected to evaluate multiple different signatures that are overlapping. The use of both scRNAseq- (Cluster 11) and snRNAseq- (Microglia 13) derived signatures help to address the concern that the type of single cell preparation (living cell vs. nucleus isolation) could influence the nature of the signature.

Moreover, while RNA signatures provide a very useful entry into the characterization of a putative cell subtype or state, they are limited in their ability to guide further functional studies; therefore, the next generation of model systems will require validation at the protein level. This is currently limited by the availability of human datasets defining microglial subsets at the protein level. Second, a signature may define a cell state, but it can be composed of distinct transcriptional programs that co-occur in a certain context. This is well-described for response to Type 1 interferons^24^, in which at least 5 transcriptional programs can be resolved. Our data suggests that this is also the case for the DAM signatures, as in the context of two different model systems (HMC3 and iMG) and Entinostat or Vorinostat exposure - illustrated in **Figure 3A** - there are at least two distinct subgroups of genes present in each of the signatures. The HDAC inhibitors engage one of these and appear to reduce expression of the other, small subgroup of genes that appears to be upregulated in the baseline state of these *in vitro* model systems in which microglia-like cells are likely to be somewhat activated. Further work will be needed to understand the role of each of these two gene subgroups. The larger subgroup, which is expressed at higher levels following exposure to HDAC inhibitors contains most of the key markers that the community has used to define the DAM signatures. This subgroup also contains *MITF* which is a transcriptional factor that has been recently proposed as a regulator of the iMG Cluster 2+8^9^. MITF expression is enhanced in both of our model systems (HMC3 microglia and iMG) following HDAC inhibitor treatment (**Figure 4D**). The authors of this study further report *ABCA1*, *APOE*, *GPNMB*, *LPL* and *TREM2* as key markers that overlap between their iPSC-derived DAM model and human brain biopsy samples when integrated into their dataset^25^. Interestingly, in our model systems, we also observe a significant increase of *ABCA1*, *APOE*, *LPL*, further confirming their data (**Table S3**). The third challenge of studying RNA-defined cell subsets involves the relation of RNA signatures, which are very dynamic, to cellular function. Our data illustrate this in that Entinostat appears to have a very strong effect on the RNA signatures when compared to Vorinostat, but Vorinostat has the stronger effect when it comes to Dextran and Aβ1-42 uptake (**Figure 5A**). This illustrates the limitation of RNA-based signatures in studying cellular functions such as Aβ1-42 uptake that may be the more clinically relevant outcome measure. Nonetheless, the DAM signatures were critical to the prioritization of these tool compounds.

The current literature on human DAM or DAM-like states does not provide any comprehensive information on whether microglia have to transition through the DAM1 state in order to become DAM2. In fact, until recently, it was not clear whether DAM or DAM-like states existed among human microglia^5,26,27^. Driven by technological advances, novel, emerging datasets all support the existence of DAM-like states in humans^6,10,25,28,29^. In our analysis of our sc- and snRNAseq datasets with regards to DAM1 and DAM2 signature expression, we observed microglia with DAM2-specific expression to be focused to regions that also showed a high expression in DAM1 marker genes, suggesting that DAM2 might arise from DAM1, but that not all microglia might transition from a DAM1 to a DAM2 state. In fact, the average proportion of DAM2 microglia is ∼1%^6^ of all microglia in older individuals (**Figure 2C-D**). On the other hand, as DAM1 marker expression is spread across almost half of the Tuddenham *et al*. ^6^ dataset, DAM1 may represent an aging- or senescence- associated microglial cluster, supported by the high expression of *SPP1* and *APOE* which also serve as markers for senescent microglia^30,31^. The live primary human microglia profiled in this study also undergo more manipulation (the effect of which is minimized by keeping the experimental pipeline on ice) than the nuclei derived from flash-frozen tissue, and this may contribute to some of the observed DAM1 changes. Similarly, whether DAM represent a subtype of microglia or rather state of reaction as result of a changed microenvironment remains to be further defined and discussed by the community^32^.

Our *in silico* analyses (**Figure 2B, Figure S2A**) prioritized a broad range of compounds with HDAC inhibition properties. Our subsequent transcriptomic and functional studies following Vorinostat and Entinostat exposure have validated this initial observation. This report joins a growing literature implicating HDAC activity in neurodegeneration and in microglial function in particular. For example, HDAC inhibition has been suggested to perturb the state of microglial activation^33^ and to cause functional changes, including suppression of cytokine and chemokine^34^ secretion. Further, ablation of HDAC1/HDAC2 in mice is reported to enhance microglial amyloid phagocytosis and to decrease amyloid load in an amyloid proteinopathy model^35^. Further, HDAC2 is implicated in the negative regulation of memory and synaptic plasticity and has been reported to be increased in postmortem samples from AD patients^36,37^. Similarly, HDAC6 has been reported to be increased in postmortem samples from individuals with AD and may be involved in metabolism of tau^38^.

Aside from enhanced Aβ 1-42 uptake which are consistent with previous reports on the MITF-dependent DAM model that has enhanced phagocytosis^9^, our functional analyses also yielded a specific downregulation of TNFα- or IFNy-induced MCP-1 secretion in both Vorinostat- and Entinostat- treated cells. MCP-1 has emerged as a cytokine with pivotal roles in many CNS disorders: it is present in senile plaques and reactive microglia in AD (summarized in ^39^). Moreover, elevated MCP-1 serum levels are increased in mild cognitive impairment (MCI) as well as in mild AD ^40^. Additionally, in cerebrospinal fluid (CSF), MCP-1 levels are significantly increased and positively correlated with ptau and ß-amyloid levels ^41^. Thus, microglia enriched in DAM-related signatures may be having multiple effects in AD, reducing Aβ1-42 load and possibly reducing leukocyte recruitment, rather implying a beneficial than detrimental role in neurodegeneration; however, further studies are needed to address these hypotheses.

The development of model systems with which to study human DAM physiology and function bears great potential. Our understanding of potentially progressive states of DAM, their role in disease as well as the expression of specific markers in these microglial subsets holds great potential for the development of therapeutic strategies with which to tackle neurodegenerative disease. Once our understanding of whether DAM play a detrimental or rather beneficial role at a certain stage in disease, is established, strategies for modulating them can be tested to either increase or decrease their numbers in a given condition.

## Supporting information

Table S1_Signatures

Table S2_CMAP Output

Table S3A_HMC3 results

Table S3B_iMG results

Table S4_DEGs

## Acknowledgements

The work was supported by the Chan-Zuckerberg Initiative’s Neurodegeneration Challenge Network grant CS-02018-191971. Some of the work also emerged from support from NIH/NIA grants R01 AG070438, U01 AG061356, RF1 AG057473, R01AG048015. Research reported in this publication was supported by the National Institute of General Medical Sciences of the National Institutes of Health under Award Number T32GM007367 and by the National Cancer Institute of the National Institutes of Health under Award Number F30CA261090. AAS is supported by The Thompson Foundation (TAME-AD) and the Henry and Marilyn Taub Foundation.

Research reported in this publication was partially performed in the Columbia Center for Translational Immunology and P&S Flow Cytometry Core and the Sulzberger Genome Center at Columbia University. This research was funded in part through the NIH/NCI Cancer Center Support Grant P30CA013696 and used the Genomics and High Throughput Screening Shared Resource.

All illustrations were created with BioRender.com

## Material & Methods

### Analysis of DAM1 and DAM2 marker gene expression in previously published single-cell and single-nucleus human microglial datasets

To plot DAM markers across the single-cell^6^ and single-nucleus datasets^10^, the schex package^42^ was used to group cells into hexagonal bins to ensure that expression of overlapping cells could be better visualized. Bin number was chosen to have relatively uniform numbers of cells across the data. For the single-cell microglial data^6^, the median number of cells per hexagon was 25, for the single-nucleus data the median number of cells per hexagon was 22. Genes were plotted across the hex-binned UMAP space, with color corresponding to log-normalized gene expression score. Similarly, module scores for the top 10 DAM1 genes and top 20 DAM2 genes were plotted across the hex-binned UMAP for both single-cell and single-nucleus microglial data. DAM1 (top10 genes): A*POE, H2-D1, B2M, FTH1, CSTB, LYZ2, CTSB, TYROBP, TIMP2, CTSD.* DAM2 (top20 genes): *ANK, CD9, CD65, SERINE2, SPP1, CADM1, CD68, CTSZ, AXL, CLEC7A, CTSA, CD52, CSF1, CCL6, LPL, CTSL, CST7, ITGAX, GUSB, HIF1A*.

Leveraging the Connectivity Map to identify pharmacological targets for *in vitro* recapitulation of the identified DAM clusters in single-cell and single-nucleus human microglial data The Connectivity Map (CMAP) is a catalog of gene expression signatures for a series of different genetic and pharmacologic perturbations across a wide variety of human cell lines ^12^

To identify chemical targets that might drive signatures associated with the signature of our identified Disease-associated microglial cluster *in vitro*, upregulated gene lists for this cluster were assembled corresponding to genes upregulated in comparison to three or more clusters from our previously published sc-RNA Seq data of human microglia ^6^. Additionally, we also performed predictions on a DAM-like cluster from a previously published single-nucleus RNA-seq dataset ^10^. This subcluster exhibited strong transcriptional overlap with Cluster 11 reported in Tuddenham et al. ^6^ and it was therefore chosen to examine both the predicted regulators for the upregulated signature for the single-nucleus cluster as well as predictions on the overlapping gene set between these two clusters with the aim to refine our predictions to drugs that would target the most crucial core regulatory signature of the DAM-like signature in humans.

The web interface found at clue.io, was used to interface with the CMAP database, and the *ListMaker* tool was used to assemble lists which were then submitted as inputs to the *Query* tool. The version 1.0 L1000 gene expression data compendium was used for all analyses. Output lists were downloaded and ranked by “median_tau_score”. Chemical perturbagens of interest were selected from those with a “median_tau_score” above 90 and chosen based on prior knowledge and the pathways they targeted. As compounds for further validation, compounds that came up to mimic both transcriptomic signatures, of Cluster 11 ^6^ and Microglia 13 ^10^ were selected. Full output lists from CMAP for Cluster 11 and Microglia 13 can be found in **Table S2**.

### Drug screening in the HMC3 model system to identify drugs driving the Disease-associated microglial- (DAM-) like signature

Compounds of interest were obtained from a wide range of reputable vendors (see **Table 1**) and resuspended in DMSO (Sigma-Aldrich, Cat #:472301). To keep the design of our experiment as similar as possible to the CMAP study, the target stock concentration was 10 mM. Dose titration with doses ranging from 0.01 µM to 0.1 mM was conducted to determine the highest tolerable dose for each compound. Each concentration of drug was plated in triplicate with early-passage HMC3 cells (ATCC; Cat #: CRL-3304), and cell viability was read out using using MTT assay, by incubating the cells in 0.25mg/ml MTT (invitrogen; Catalog #: M6494) in PBS (Corning, Cat #:21-040-CV) for 1hr at 37°C, following removal of MTT solution and addition of 200µl DMSO (Sigma-Aldrich, Cat #:472301) and further incubation of the cells for 15 min at 37°C before measuring the absorbance at 570 nm with a Tecan Infinite® 200 PRO plate reader (Tecan; Cat#: 30050303). An optimal dose of each drug was then chosen based on cell morphology and viability. Subsequently, 0.35x10^6^ HMC3 microglial cells were seeded into a 6-well plate and incubated o.n.. The next day, microglia were treated with the respective concentrations of Vorinostat (1µM; Ambeed; Cat #: A234507), Cholic acid (10µM; Cayman chemical; Cat #: 20250), Flavokavain B (10µM; ChromaDex; Cat #: ASB-00006058- 005), Wiskostatin (1µM; Cayman chemical; Cat #: 15047), Trimipramine (10µM; Cayman chemical; Cat #: 15921), Naftopidil (10µM; APExBIO; Cat #: 57149-07-2), Ramipril (10µM; APExBIO; Cat #: B2208), Valporic acid (90µg/ml Sigma; Cat #: PHR1061), Geranylgeraniol (10µM; Sigma; Cat #: 24034-73-9), Entinostat (10µM; Ambeed; Cat #: A122285), Amiodarone hydrochloride (10µM; Sigma; Cat #: A8423), Temozolomide (100µM; APExBIO; Cat #: B1399) or DMSO (Sigma-Aldrich, Cat #:472301) as control and incubated for 6hrs and 24hrs before harvest for RNA extraction. Lysis was performed in-well with buffer RLT (Qiagen; Cat #: 74136) containing 2-Mercaptoethanol (Thermo Fisher Scientific, Cat #:63689), and RNA extraction was performed with the Qiagen RNEasy mini plus kit (Qiagen; Cat #: 74136) following the manufacturer’s instructions. gDNA eliminator columns were used to remove contaminating genomic DNA. Initial RNA quality and quantity was assessed using Nanodrop (ThermoFisher Scientific) followed by cDNA preparation using the BioRad iScript cDNA Synthesis kit (BioRad; Cat #:1708891). cDNA was subsequently purified with AMPure XP beads (Thermo Fisher Scientific; Cat #: A63880) using a 1:1.8 ratio of cDNA: beads, concentration determined using Qubit HS DNA quantification kit (Thermo Fisher Scientific) and subsequently adjusted to 1ng/µl.

### Quantitative RT-PCR analysis

Quantitative real-time PCR reactions to amplify 1 ng of total cDNA were performed in a QuantStudio^TM^ 3 Real-Time PCR Cycler (A28132, Applied Biosystems) using the Applied Biosystems Fast SYBR Green Master Mix (Thermo Fisher Scientific; Cat #: 4385612). CT values were normalized using Hypoxanthine Phosphoribosyltransferase 1 (*Hprt1*) as the housekeeping gene. Primers were tested for their efficiency beforehand, and the ΔΔCt-method was applied for analysis of relative gene expression. The melting curves of each product were analyzed to ensure the specificity of the PCR product. The following primers were used: ***HPRT1*** - fw: CCTGGCGTCGTGATTAGTGAT, rev: AGACGTTCAGTCCTGTCCATAA; ***CD9*** – fw: GTT TCT TGC TCG AAG ATG CTC, rev: CAC CAA GTG CAT CAA ATA CCT G ; ***CTSD*** - fw: CTTCGACAACCTGATGCAGC, rev: TACTTGGAGTCTGTGCCACC; ***SPP1*** - fw: GCCGAGGTGATAGTGTGGTT, rev: AACGGGGATGGCCTTGTATG. For visualization, the mean for each gene is shown with error bars that denote standard deviation. Individual points are plotted to visualize the distribution of the data. For statistical analysis, log-fold change values in comparison to DMSO-treated control samples were analyzed using Kruskal Wallis test followed by Dunn’s multiple comparison test. Data was subsequently plotted in GraphPad Prism 9.2.0.

### Bulk RNA-Seq of HMC3 microglia treated with drugs driving the Disease-associated microglial (DAM-) signature

In brief, 0.75-1x10^6^ HMC3 microglial cells were seeded into a 6.5cm or 10cm dish and incubated o.n. The next day, microglia were treated with the respective concentrations of Vorinostat (1µM; Ambeed; Cat #: A234507), Entinostat (10µM; Ambeed; Cat #: A122285) or DMSO (Sigma-Aldrich, Cat #:472301) as control and incubated for 24 hrs before harvest. Cells were trypsinized (Gen Clone; Cat #:25-510F), counted, the cell viability was assessed. Cells were then washed with ice-cold PBS (Corning, Cat #:21-040-CV) and resuspended in 350μl RLT Lysis buffer (Qiagen; Cat #: 74136) containing 2-Mercaptoethanol (Thermo Fisher Scientific, Cat #:63689), and isolated using Qiagen Plus Mini Kit (Qiagen; Cat #: 74136). RNA quality was assessed using the TapeStation (Agilent) prior to further processing for RNA sequencing. mRNA libraries were prepped using poly-A pull-down to enrich mRNAs from total RNA samples followed by Illumina TruSeq Stranded mRNA Library prep (Illumina, Cat#: 20020595), in accordance with manufacturer recommendations, and using IDT for Illumina TruSeq DNA UD Indices (Illumina, Cat#: 20022370) for adapters. Briefly, 500ng of total RNA was used for purification and fragmentation of mRNA. Purified mRNA underwent first and second strand cDNA synthesis. cDNA was then adenylated, ligated to Illumina sequencing adapters, and amplified by PCR (using 10 cycles). The cDNA libraries were quantified using Fragment Analyzer 5300 (Advanced Analytical) kit FA-NGS-HS (Agilent, Cat#: DNF-474-1000) and Spectramax M2 (Molecular Devices) kit Picogreen (Life Technologies, Cat#: P7589). Libraries were sequenced on an Illumina NovaSeq6000 sequencer at Columbia Genome Center. We multiplex samples in each lane, which yields targeted number of paired-end 100bp reads for each sample. We used RTA (Illumina) for base calling and bcl2fastq2 (version 2.20) for converting BCL to fastq format, coupled with adaptor trimming. We performed a pseudoalignment to a kallisto index created from transcriptomes (Ensembl v111, Human:GRCh38.p14; Mouse:GRCm39; mRatBN7.2) using kallisto (0.50.1). The references and kallisto version were updated on April 29, 2024 to ensembl v111 and kallisto 0.50.1.

### hiPSC maintenance and differentiation into microglia-like cells (iMG) and treatment with DAM-inducing compounds

hiPSCs were maintained in StemFlex media (ThermoFisher) on reduced growth factor Cultrex BME (Biotechne, Cat.# 3434-010-02), and routinely split 1-2 times a week with ReLesSR™ (Stem Cell Technologies) without ROCKi as described previously ^43^. hiPSCs were differentiated into iMGLs as described previously with minor adaptations ^44,45^. In brief, FA000010 (RUCDR/Infinity BiologyX) iPSCs were differentiated into hematopoietic precursors cells (HPCs) using the STEMdiff Hematopoietic kit (STEMCELL Technologies) largely by manufacturer’s instructions. In brief, on day -1 iPSCs were detached with ReLeSR and passaged to achieve a density of 1–2 aggregates/cm^2^ of 100-150 cells. Multiple densities were plated in parallel. On day 0, colonies of appropriate density were switched to Medium A from the STEMdiff Hematopoietic Kit to initiate HPC differentiation. On day 3, cells were switched to Medium B with a full media change and fed again with a full media change on day 5. Cells remained in Medium B for the rest of the HPC differentiation period with Medium B overlay feeds every other day. HPCs were collected 3 independent times by gently removing the floating population with a serological pipette at days 11, 13 and 15 (or days 12, 14 and 16).

HPCs were either cryobanked in 45% Medium B, 45% knockout serum replacement (ThermoFisher) and 10% DMSO and stored in liquid nitrogen or directly plated for iMGL induction. HPCs were terminally differentiated at 28,000-35,000 cells/cm^2^ in microglia medium (DMEM/F12, 2X insulin-transferrin-selenite, 2X B27, 0.5X N2, 1X glutamax, 1X non-essential amino acids, 400 mM monothioglycerol, and 5 mg/mL human insulin (ThermoFisher)) freshly supplemented with 100 ng/mL IL-34, 25 ng/mL M-CSF (R&D System) and 50 ng/mL TGFβ1 (STEMCELL Technologies) for every other day until day 24. On day 25, 100 ng/mL CD200 (Bon Opus Biosciences) and 100 ng/mL CX3CL1 (R&D Systems) were added to Microglia medium to mimic a brain-like environment. Differentiated microglia were cultured in 500µl microglia medium (DMEM/F12, 2X insulin-transferrin-selenite, 2X B27, 0.5X N2, 1X Glutamax, 1X non-essential amino acids, 400 mM monothioglycerol, and 5 mg/mL human insulin (ThermoFisher)). For compound treatment on day 28/29 of differentiation post HPC, 2x solutions were prepared for Vorinostat (final concentration: 0.1µM; Ambeed; Cat#: A234507) or DMSO as control and subsequently 500µl of 2x solutions was added to each well and incubated for 24 hrs. Cells were subsequently harvested by pipetting up and down using PBS, counted and the cell viability was assessed. Cells were then washed with ice-cold PBS (Corning, Cat #:21-040-CV), resuspended in 350μl RLT Lysis buffer (Qiagen; Cat #: 74136) containing 2-Mercaptoethanol (Thermo Fisher Scientific, Cat #:63689) and stored until further processing.

### Bulk RNA-Seq of DAM-treated iMGs

Subsequently, RNA was isolated using Qiagen Plus Mini Kit (Qiagen; Cat #: 74136). RNA quality was assessed using TapeStation (Agilent) prior to further processing for RNA sequencing. mRNA libraries were prepped using poly-A pull-down to enrich mRNAs from total RNA samples followed by Illumina TruSeq Stranded mRNA Library prep (Illumina, Cat#: 20020595), in accordance with manufacturer recommendations, and using IDT for Illumina TruSeq DNA UD Indices (Illumina, Cat#: 20022370) for adapters. Briefly, 500ng of total RNA was used for purification and fragmentation of mRNA. Purified mRNA underwent first and second strand cDNA synthesis, the final PCR step has been modified using the KAPA HiFi HotStart Ready Mix.The cDNA libraries were quantified using Fragment Analyzer 5300 (Advanced Analytical) kit FA-NGS-HS (Agilent, Cat#: DNF-474-1000) and Spectramax M2 (Molecular Devices) kit Picogreen (Life Technologies, Cat#: P7589). Libraries are then sequenced using Element AVITI at Columbia Genome Center. We multiplex samples in each lane, which yields targeted number of paired-end 75bp reads reads for each sample. We use bases2fastq version 1.7.0.1196148384 for converting BCL to fastq format, coupled with adaptor trimming. We perform a pseudoalignment to a kallisto index created from transcriptomes (Ensembl v111, Human:GRCh38.p14; Mouse:GRCm39; mRatBN7.2) using kallisto (0.50.1). The references and kallisto version were updated on April 29, 2024 to ensembl v111 and kallisto 0.50.1.

### Data pre-processing and analysis of bulk RNA Sequencing of compound-treated HMC3 and iMG DAM-models

Illumina TruSeq chemistry was used for library construction after poly-A pull-down was used to augment mRNA from total RNA Samples. Illumina NovaSeq 6000 was used at the Columbia Genome Center to sequence libraries. Samples in each lane were multiplexed, generating the selected number of paired-end 100bp reads for each sample. For base calling and converting BCL to fastq format, RTA (Illumina) and bcl2fastq2 (version 2.20) were used in conjunction with adaptor trimming, respectively. Pseudoalignment to a kallisto index created from transcriptomes (Ensembl v111, Human:GRCh38.p14) using kallisto (0.50.1).

The DESeq2^18^ package implemented within R (4.4.1) was used to test for differentially expressed genes between treatment conditions. By creating a DESeq object, count values were compared across treatments and controls using generalized linear models in single factor model (treatment). To identify any outlying samples, variance stabilizing transformation was used on the same DEseq object, and PCA was performed. For the purposes of visualizing expression, heatmaps were generated using the TPM metric of expression. For volcano plots, genes with a baseline mean count > 100 or genes within the specified gene list were included. To determine significant upregulation in any particular gene set, a Wald test and its statistic were used to generate a p-value and a resulting Benjamini-Hochberg (FDR) value, where each gene set was analyzed independently of the others. After applying this test, any gene with a positive log2 fold-change and an FDR < 0.05 was considered to have statistically significant upregulation.

### Phagocytosis Assays

#### Aß Phagocytosis as a proxy for phagocytic behavior in a neurodegenerative context

0.1mg human beta-Amyloid (1-42), HiLyte™ Fluor 647-labeled (Anaspec Inc., Cat#: AS- 64161) was resuspended to a 4µM stock in 50µl 1% Na4OH and 3950µl Corning™ Eagle’s Minimum Essential Medium (MEM) (Corning, Cat#: 10-009-CV) and aliquots stored at -80°C until usage. For phagocytosis assay, 75K HMC3 microglia were seeded in a 24-well plate and incubated at 37°C, 5% CO2 o.n. The next day, cells were treated with Vorinostat (1µM;Ambeed; Cat #: A234507), Entinostat (10µM; Ambeed; Cat #: A122285) and DMSO (Sigma-Aldrich, Cat #:472301) as control and incubated for another 24hrs. Media was removed and cells were subsequently incubated in 50nM Aß-containing complete DMEM media (10% FCS (Fisher Scientific, Cat#: 10-438-026), 1% P/S (Gibco, Cat#: 15140-122) or DMSO as a control, additionally containing either 5µM Cytochalasin D (Sigma-Aldrich, Cat#: C8273) as a negative control for phagocytosis, or DMSO as control for Cytochalasin D treatment. Cells were incubated for 1hr at 37°, 5% CO2 and subsequently washed twice with pre-warmed PBS (Corning, Cat #:21-040-CV) before harvest using Trypsin (Gen Clone; Cat #:25-510F). Cells were subsequently harvested in Flow tubes (MTC Bio, Cat#: T9005), centrifuged at 1500 rpm for 5 min at 4°C and resuspended in 500µl Cell Staining buffer (BioLegend, Cat#: 420201) containing 1:4000 dilution of SYTOX Blue (ThermoFisher, Cat#: S34857) for labeling of live/dead cells. To assess phagocytosis, samples were subsequently assessed using Aurora 3L analyzer (Cytek Bio), followed by analysis using FlowJo_v10.8.1. Cells were gated for single cells, live cells and then Alexa647-positive cells and analyzed percentages subsequently normalized to DMSO control for each experiment to allow comparison across experiments performed on different days. For statistical analysis, log-fold change values in comparison to DMSO-treated control samples were analyzed using one-way ANOVA followed by Dunnett’s multiple comparison test. Data was subsequently plotted in GraphPad Prism 9.2.0.

#### Phagocytosis of pH-rhodo Dextran as a proxy for Macropinocytosis

pHrhodo Green Dextran (life technologies, Cat#: P10361) was resuspended in 0.5ml deionized water (Sigma-Aldrich, Cat#: 95284) to 1mg/ml stock solution and aliquots stored at -20°C. For phagocytosis assay, 75K HMC3 microglia were seeded in a 24-well plate and incubated at 37°C, 5% CO2 o.n. The next day, cells were treated with Vorinostat (1µM;Ambeed; Cat #: A234507), Entinostat (10µM; Ambeed; Cat #: A122285) and DMSO control (Sigma-Aldrich, Cat #:472301) as control and incubated for another 24 hrs. Media was removed and cells were subsequently incubated in 100µg/ml pHrhodo Green Dextran-containing complete DMEM media (10% FCS (Fisher Scientific, Cat#: 10-438-026), 1% P/S (Gibco, Cat#: 15140- 122)) or DMSO as a control, additionally containing either 5µM Cytochalasin D (Sigma-Aldrich, Cat#: C8273) as a negative control for phagocytosis, or DMSO as control for Cytochalasin D treatment. Cells were incubated for 1hr at 37°, 5% CO2 and subsequently washed twice with pre-warmed PBS (Corning, Cat #:21-040-CV) before harvest using Trypsin (Gen Clone; Cat #:25-510F) . Cells were subsequently harvested in Flow tubes (MTC Bio, Cat#: T9005), centrifuged at 1500 rpm for 5 min at 4°C and resuspended in 500µl Cell Staining buffer (BioLegend, Cat#: 420201) containing 1:4000 dilution of SYTOX Blue (ThermoFisher, Cat#: S34857) for labeling of live/dead cells. To assess phagocytosis, samples were subsequently assessed using Aurora 3L analyzer (Cytek Bio), followed by analysis using FlowJo_v10.8.1. Cells were gated for single cells, live cells and then Alexa647-positive cells and analyzed percentages subsequently normalized to DMSO control for each experiment to allow comparison across experiments performed on different days. For statistical analysis, log-fold change values in comparison to DMSO-treated control samples were analyzed using one-way ANOVA followed by Dunnett’s multiple comparison test. Data was subsequently plotted in GraphPad Prism 9.2.0.

#### Phagocytosis of pH-rhodo E.coli as a proxy for acute neuroinflammation

pHrodo™ Green *E. coli* BioParticles™ Conjugate for Phagocytosis (ThermoFisher Scientific,Cat#: P35366) were resuspended in 2 ml PBS (Corning, Cat #:21-040-CV) and incubated for 45 min in a sonicator bath to generate 1mg/ml stock suspension. For phagocytosis assay, 75K HMC3 microglia were seeded in a 24-well plate and incubated at 37°C, 5% CO2 o.n. The next day, cells were treated with Vorinostat (1µM;Ambeed; Cat #: A234507), Entinostat (10µM; Ambeed; Cat #: A122285) and DMSO (Sigma-Aldrich, Cat #:472301) as control and incubated for another 24 hrs. Media was removed and cells were subsequently incubated in 160µl *E.coli* solution containing either 5µM Cytochalasin D (Sigma-Aldrich, Cat#: C8273) as a negative control for phagocytosis, or DMSO as control for Cytochalasin D treatment. Cells were incubated for 1hr at 37°, 5% CO2 and subsequently washed twice with pre-warmed PBS (Corning, Cat #:21-040-CV) before harvest using Trypsin (Gen Clone; Cat #:25-510F). Cells were subsequently harvested in Flow tubes (MTC Bio, Cat#: T9005), centrifuged at 1500 rpm for 5 min at 4°C and resuspended in 500µl Cell Staining buffer (BioLegend, Cat#: 420201) containing 1:4000 dilution of SYTOX Blue (ThermoFisher, Cat#: S34857) for labeling of live/dead cells. To assess phagocytosis, samples were subsequently assessed using Aurora 3L analyzer (Cytek Bio), followed by analysis using FlowJo_v10.8.1. Cells were gated for single cells, live cells and then Alexa647-positive cells and analyzed percentages subsequently normalized to DMSO control for each experiment to allow comparison across experiments performed on different days. For statistical analysis, log-fold change values in comparison to DMSO-treated control samples were analyzed using one-way ANOVA followed by Dunnett’s multiple comparison test. Data was subsequently plotted in GraphPad Prism 9.2.0.

#### Cytokine Multiplex Assay

10K HMC3 cells from three different passages were seeded in 96-well plates containing 200µl of complete Eagle’s Minimum Essential Medium (EMEM; ATCC, Cat#: 30-2003) and incubated o.n. at 37°C, 5% CO2. The next day, cells were treated with compounds using the following concentrations (6 wells per condition): DMSO control (1:1000), Vorinostat (1µM), Entinostat (10µM), and incubated for 24 hrs. The next day, 2 wells/ condition were stimulated with TNF- a (0.3 µg/mL, Peprotech, Cat#: 300-02), 2 wells/ condition were stimulated with IFN-y (0.3 µg/mL, Peprotech, Cat#: 300-01A) and as a control 2 wells/condition were stimulated with H2O as control, each dissolved in a fresh dilution of the respective drug treatment in a final volume of 200µl. For subsequent cytokine expression analysis, cell culture supernatant was harvested after 10 hrs and 24 hrs and stored at -20°C until further analysis. For cytokine multiplex assay, 100µl per sample was sent to Eve Technologies (Calgary, Alberta) and analyzed using Human Cytokine Proinflammatory Focused 15-Plex Discovery Assay® Array (HDF15). Data was subsequently plotted in GraphPad Prism 9.2.0.

## Authors contributions

Conceptualization, P.L.D., V.H. and J.F.T.; Methodology: P.L.D., V.H. and J.F.T.; Investigation: V.H., J.F.T., A.B., F.G., R.P., N.C.-L., R.K.S., P.L.D.; Data Curation: V.H., J.F.T., A.B., C.C.W., N.C.-L., R.K.S., P.S., V.M., P.L.D.; Formal Analysis: V.H., J.F.T., A.B., C.C.W., N.C.-L., V.M., V.M., P.L.D.; Writing - Original Draft: V.H., P.L.D.; Funding Acquisition: P.L.D; Resources: A.S.S., P.S., V.M., P.L.D.; Supervision: P.L.D., V.H.

## Supplemental Tables

Table S1: Complete list of all gene signatures relevant to the manuscript.

Table S2: Full output lists from CMAP12 analysis for Cluster 116 and Microglia 1310.

Table S3A: Results for all inquired gene signatures in compound-treated HMC3 microglia. Table S3B: Results for all inquired gene signatures in compound-treated iPSC-derived microglia.

Table S4: Full list of differentially expressed genes between the different treatment conditions for HMC3 microglia and iPSC-derived microglia.

