## Supplementary Material for "HDAC Inhibitors recapitulate Human Disease-Associated Microglia Signatures *in vitro*"

**Supplementary Figure 1**

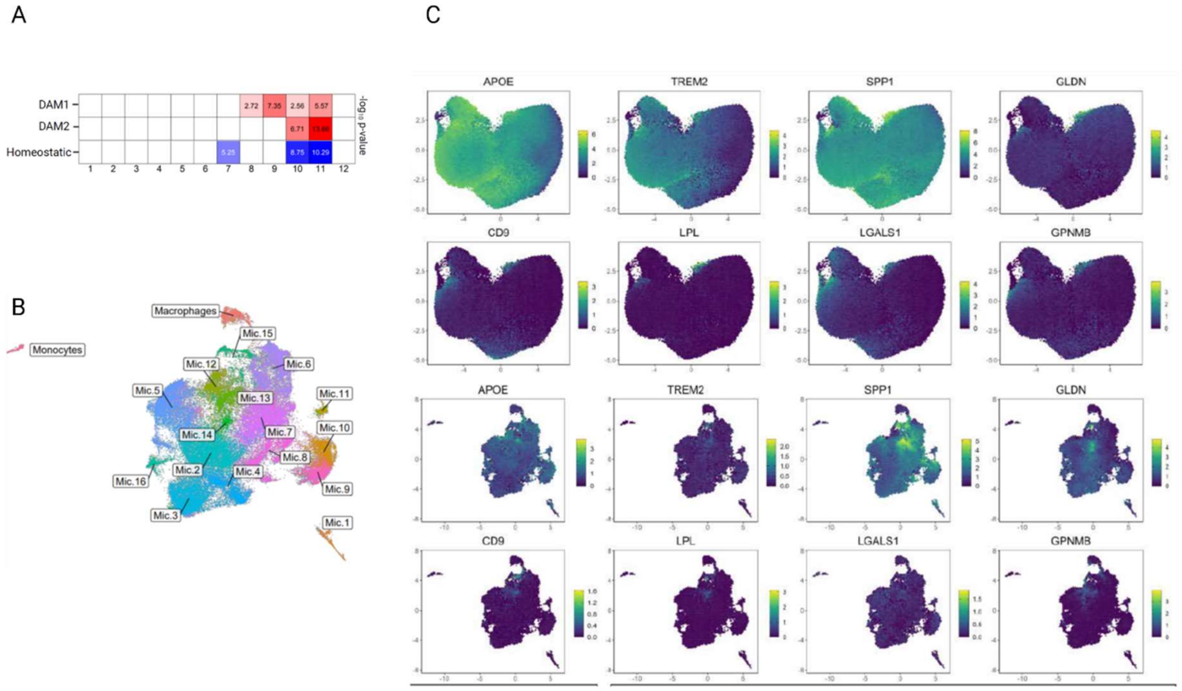


**Supplementary Figure 1. A. Statistical analysis of DAM1 and DAM2 marker enrichment analysis in microglial clusters defined in 6.** B. UMAP derived from 10, including the monocyte cluster. C. UMAP plots showing expression of DAM1 and DAM2 marker genes in human single-cell and single-nucleus microglia datasets. Enrichment of the top 10 genes for the DAM1 signature or the top 20 genes for the DAM2 signature from the original publication was calculated on a per-cell basis. Module scores were computed compared to background genes with similar levels of expression. Individual cells are colored by log-fold change of the gene set. Module scores were plotted on hex-binned UMAPs. Individual hexagons are aggregates of 50 cells on average, the plotted score per hexagon is the mean of the score across all cells aggregated within each hexagon. Scores are log-normalized counts, as shown on the color gradient bar. Yellow represents the maximal expressed value, while purple represents the lowest expression values. Selected DAM1 and DAM2 marker genes were plotted across microglial clusters from the single-cell dataset by Tuddenham et al. 2022 (upper row; 6) and the single-nucleus RNA-Seq dataset (lower row; 10).

**Supplementary Figure 2**

**A** predicted DAM compounds

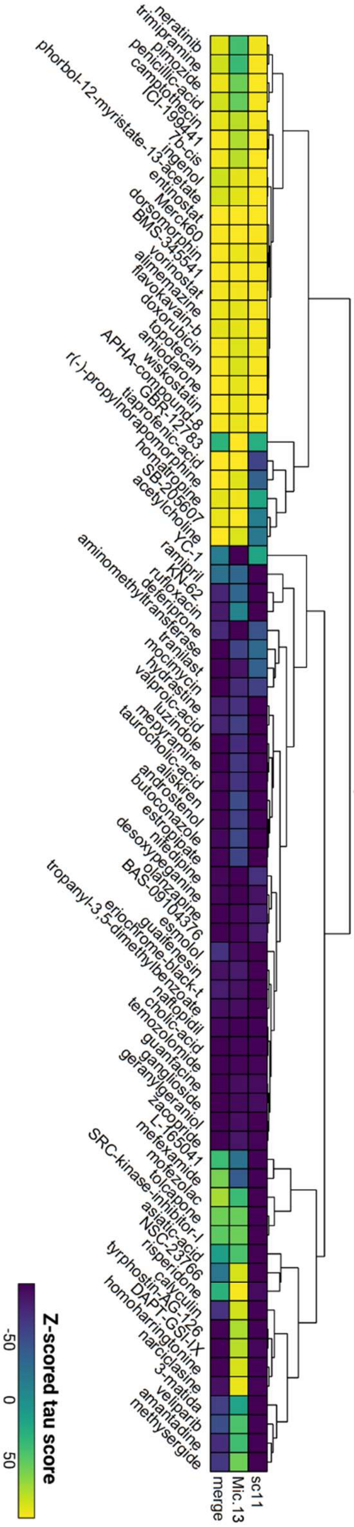

**B**

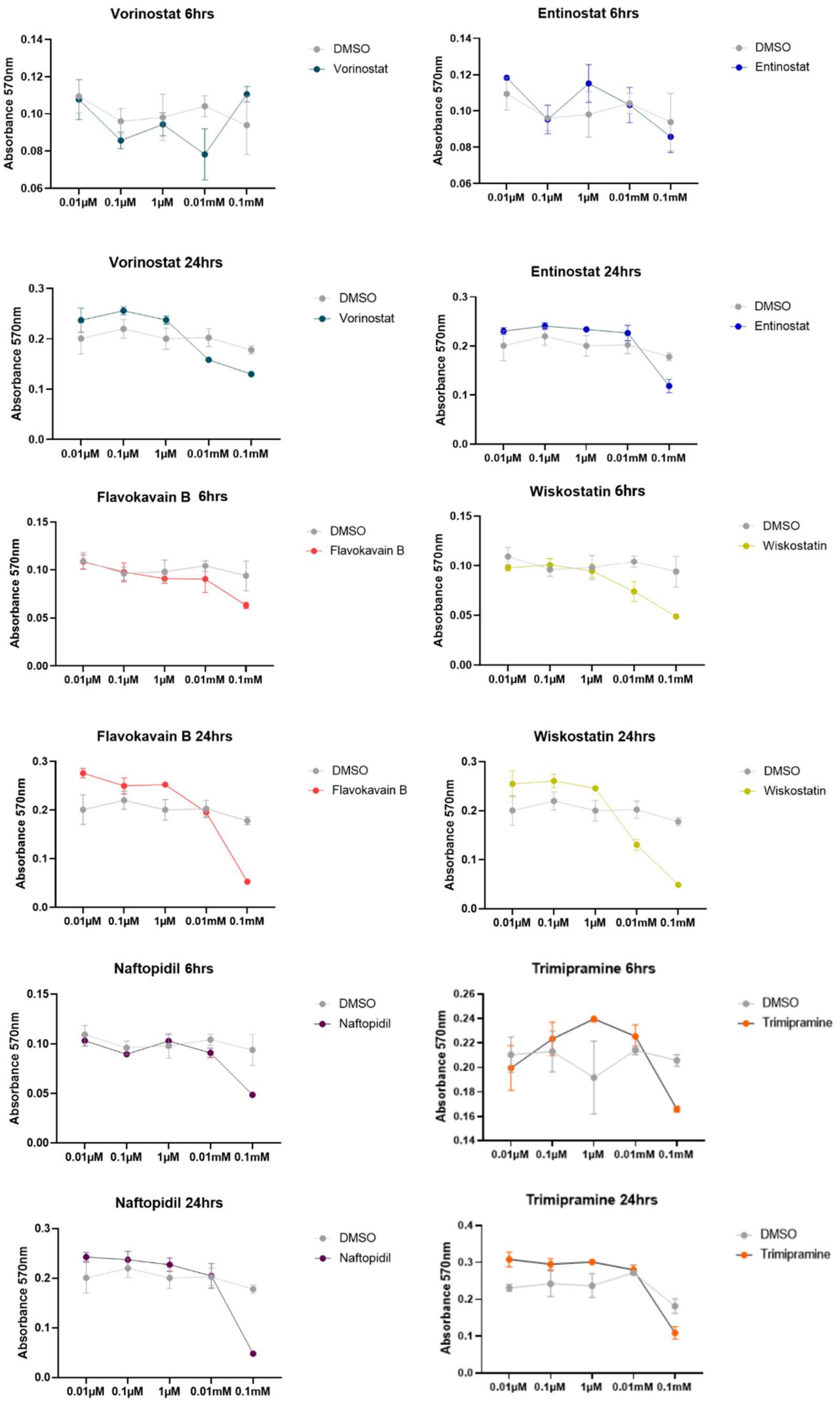
Vorinostat 6hrs
Absorbance 570nm
0.14
0.12
0.10
0.08
0.06
0.01μM 0.1μM 1μM 0.01mM 0.1mM
DMSO
Vorinostat
Entinostat 6hrs
Absorbance 570nm
0.14
0.12
0.10
0.08
0.06
0.01μM 0.1μM 1μM 0.01mM 0.1mM
DMSO
Entinostat
Vorinostat 24hrs
Absorbance 570nm
0.3
0.2
0.1
0.0
0.01μM 0.1μM 1μM 0.01mM 0.1mM
DMSO
Vorinostat
Entinostat 24hrs
Absorbance 570nm
0.3
0.2
0.1
0.0
0.01μM 0.1μM 1μM 0.01mM 0.1mM
DMSO
Entinostat
Flavokavain B 6hrs
Absorbance 570nm
0.15
0.10
0.05
0.00
0.01μM 0.1μM 1μM 0.01mM 0.1mM
DMSO
Flavokavain B
Wiskostatin 6hrs
Absorbance 570nm
0.15
0.10
0.05
0.00
0.01μM 0.1μM 1μM 0.01mM 0.1mM
DMSO
Wiskostatin
Flavokavain B 24hrs
Absorbance 570nm
0.3
0.2
0.1
0.0
0.01μM 0.1μM 1μM 0.01mM 0.1mM
DMSO
Flavokavain B
Wiskostatin 24hrs
Absorbance 570nm
0.3
0.2
0.1
0.0
0.01μM 0.1μM 1μM 0.01mM 0.1mM
DMSO
Wiskostatin
Naftopidil 6hrs
Absorbance 570nm
0.15
0.10
0.05
0.00
0.01μM 0.1μM 1μM 0.01mM 0.1mM
DMSO
Naftopidil
Trimipramine 6hrs
Absorbance 570nm
0.26
0.24
0.22
0.20
0.18
0.16
0.14
0.01μM 0.1μM 1μM 0.01mM 0.1mM
DMSO
Trimipramine
Naftopidil 24hrs
Absorbance 570nm
0.3
0.2
0.1
0.0
0.01μM 0.1μM 1μM 0.01mM 0.1mM
DMSO
Naftopidil
Trimipramine 24hrs
Absorbance 570nm
0.4
0.3
0.2
0.1
0.0
0.01μM 0.1μM 1μM 0.01mM 0.1mM
DMSO
Trimipramine

C

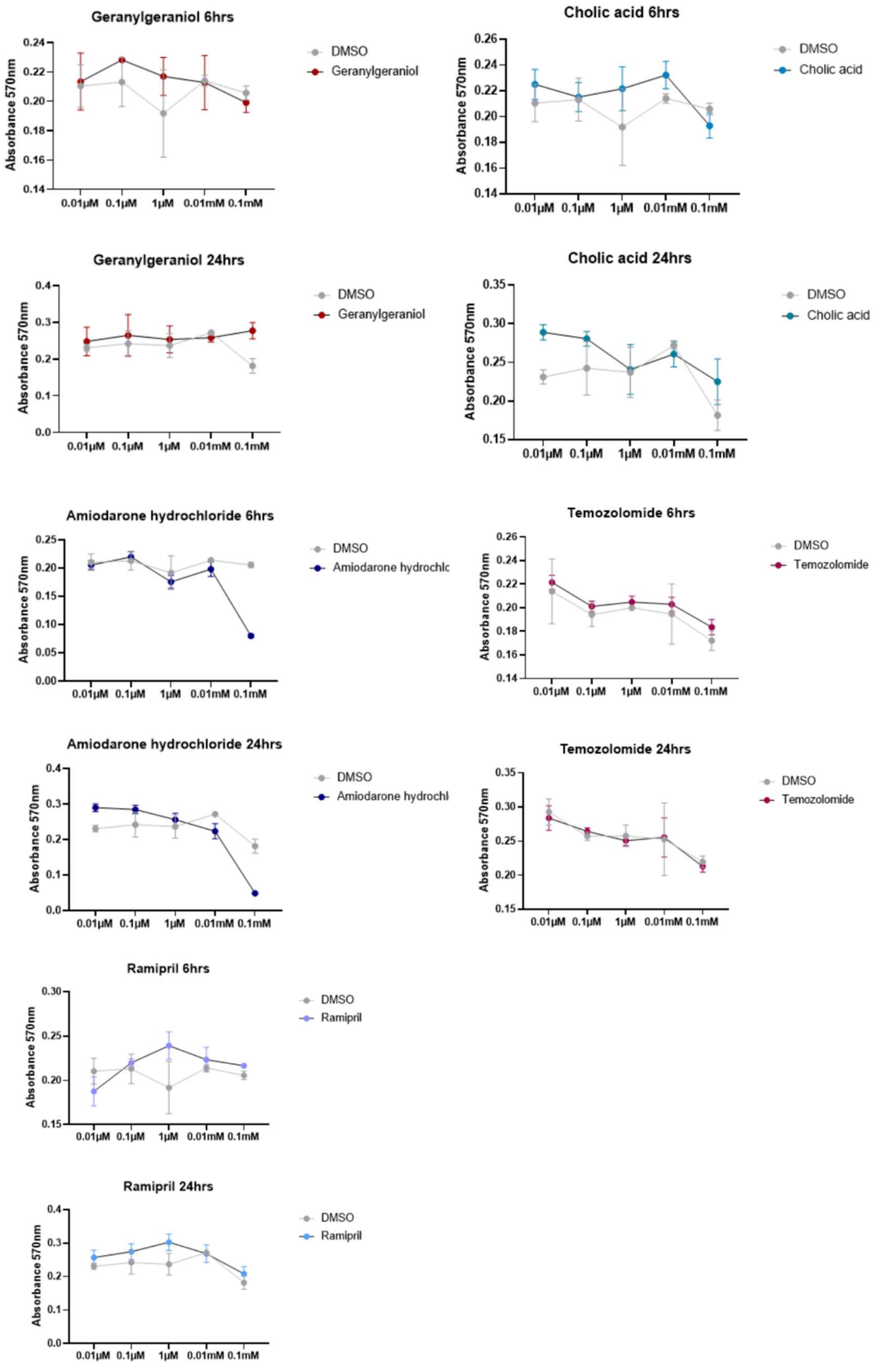
Geranylgeraniol 6hrs
Absorbance 570nm
0.24
0.22
0.20
0.18
0.16
0.14
0.01μM 0.1μM 1μM 0.01mM 0.1mM
DMSO
Geranylgeraniol
Cholic acid 6hrs
Absorbance 570nm
0.26
0.24
0.22
0.20
0.18
0.16
0.14
0.01μM 0.1μM 1μM 0.01mM 0.1mM
DMSO
Cholic acid
Geranylgeraniol 24hrs
Absorbance 570nm
0.4
0.3
0.2
0.1
0.0
0.01μM 0.1μM 1μM 0.01mM 0.1mM
DMSO
Geranylgeraniol
Cholic acid 24hrs
Absorbance 570nm
0.35
0.30
0.25
0.20
0.15
0.01μM 0.1μM 1μM 0.01mM 0.1mM
DMSO
Cholic acid
Amiodarone hydrochloride 6hrs
Absorbance 570nm
0.25
0.20
0.15
0.10
0.05
0.00
0.01μM 0.1μM 1μM 0.01mM 0.1mM
DMSO
Amiodarone hydrochlo
Temozolomide 6hrs
Absorbance 570nm
0.26
0.24
0.22
0.20
0.18
0.16
0.14
0.01μM 0.1μM 1μM 0.01mM 0.1mM
DMSO
Temozolomide
Amiodarone hydrochloride 24hrs
Absorbance 570nm
0.4
0.3
0.2
0.1
0.0
0.01μM 0.1μM 1μM 0.01mM 0.1mM
DMSO
Amiodarone hydrochl
Temozolomide 24hrs
Absorbance 570nm
0.35
0.30
0.25
0.20
0.15
0.01μM 0.1μM 1μM 0.01mM 0.1mM
DMSO
Temozolomide
Ramipril 6hrs
Absorbance 570nm
0.30
0.25
0.20
0.15
0.01μM 0.1μM 1μM 0.01mM 0.1mM
DMSO
Ramipril
Ramipril 24hrs
Absorbance 570nm
0.4
0.3
0.2
0.1
0.0
0.01μM 0.1μM 1μM 0.01mM 0.1mM
DMSO
Ramipril

**Supplementary Figure 2. A. Overview of the full list of identified compounds** to affect the expression of Cluster 11 [1] or Microglia 13 signature as a result of the *in silico* compound screen using the CMAP [2] resource. The Connectivity Map (CMAP) was used to identify compounds that are predicted to upregulate or downregulate gene sets associated with either of the DAM-like clusters from single-cell or single-nucleus data. Heatmaps depict the Z-scored tau score for each compound following query analysis. Clustering of tau scores across microglial clusters 11, 13, or the merged signature was performed with absolute linkage. **B-C. Titration experiments for validation experiments following *in silico* prediction analysis.** To determine the right dose for the treatment of HMC3 microglia, each compound (predicted upregulation: **B**; predicted downregulation: **C**) was titrated on HMC3 microglia using 5 different concentrations covering a range from 0.1mM to 0.01µM (0.1mM, 0.01mM, 1µM, 0.1µM, 0.01µM). Following 6hrs and 24hrs treatment, cell viability was assessed using MTT viability assay via fluorescent readout at 570 nm. Results for each tested compound are shown in comparison to DMSO control. Data points/concentration are derived from treatment in triplicates as mean ± SD. Compound-treatment is depicted in color, whereas DMSO controls are visualized in gray.

**Supplementary Figure 3**

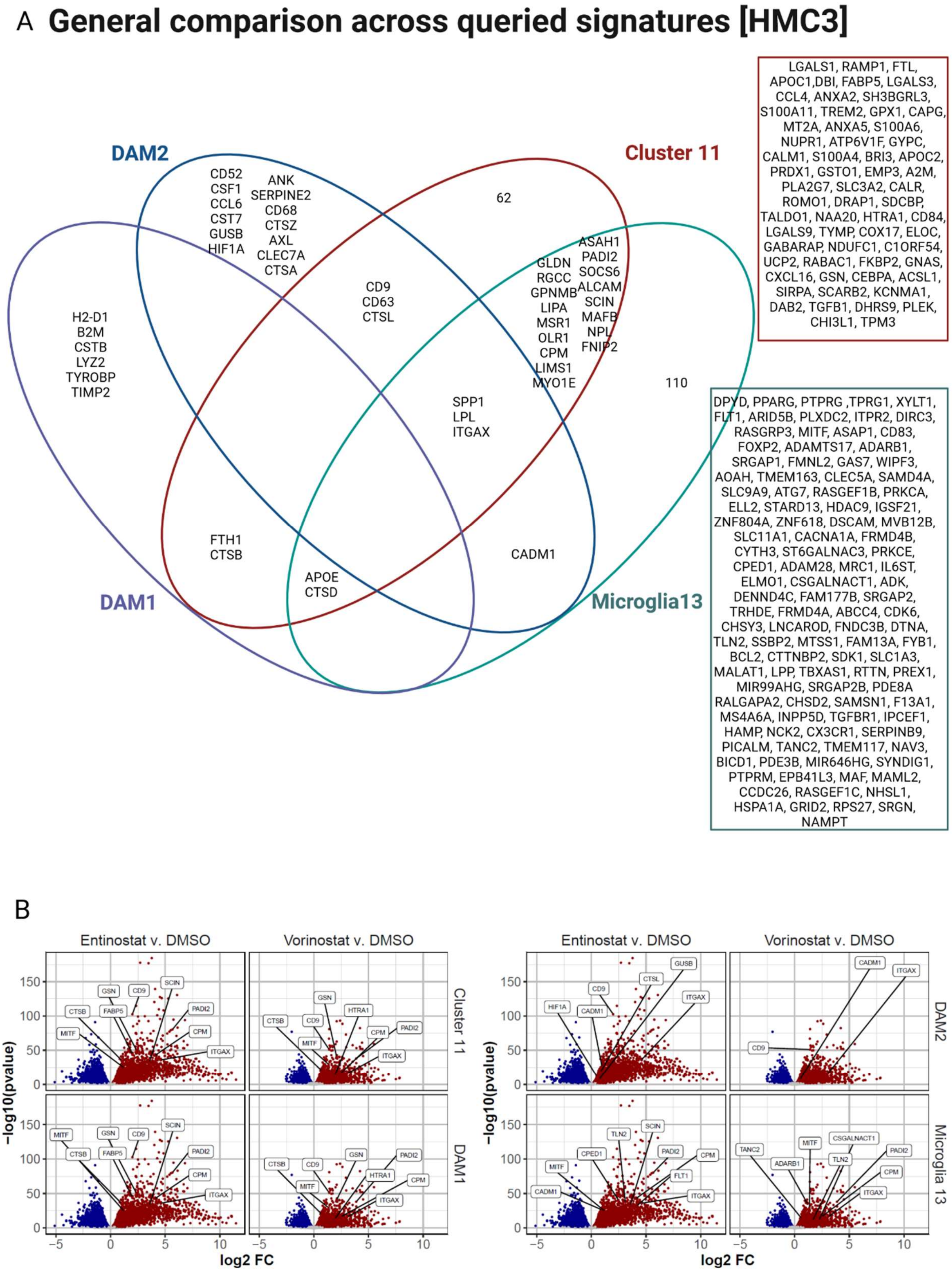

C Entinostat [HMC3]

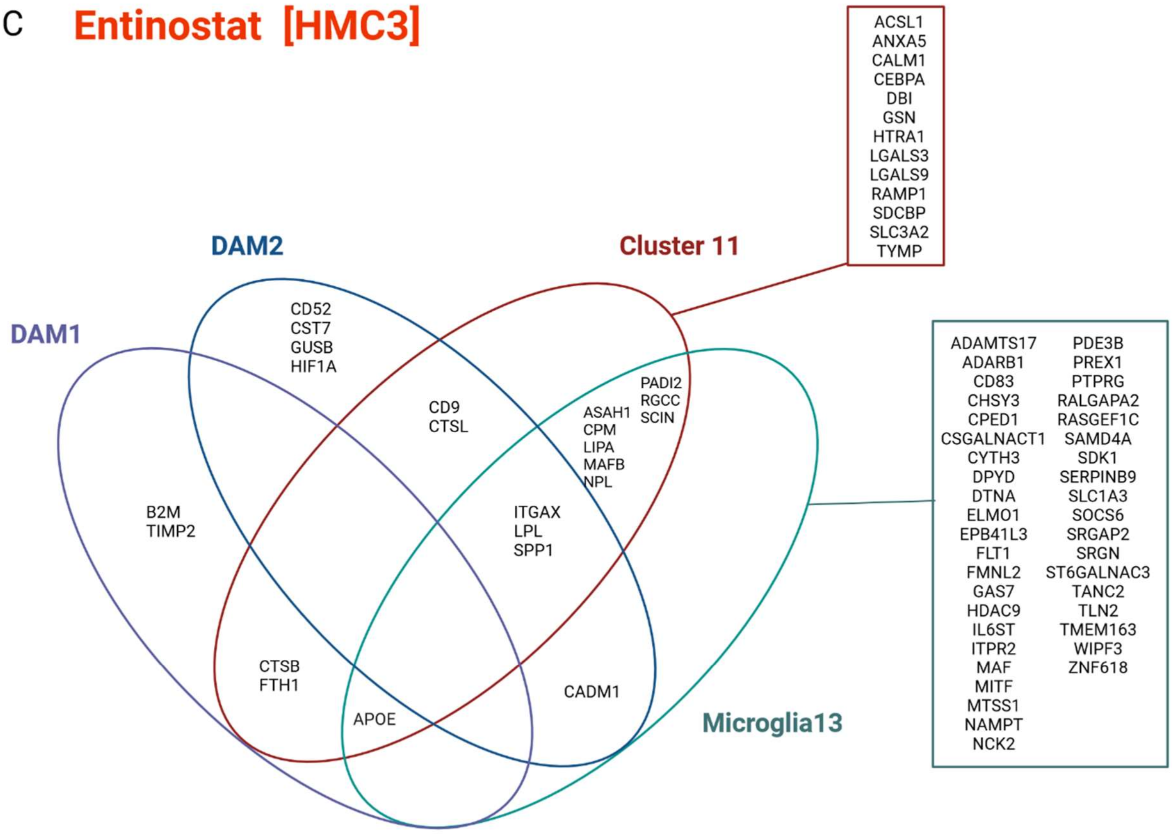


D Vorinostat [HMC3]

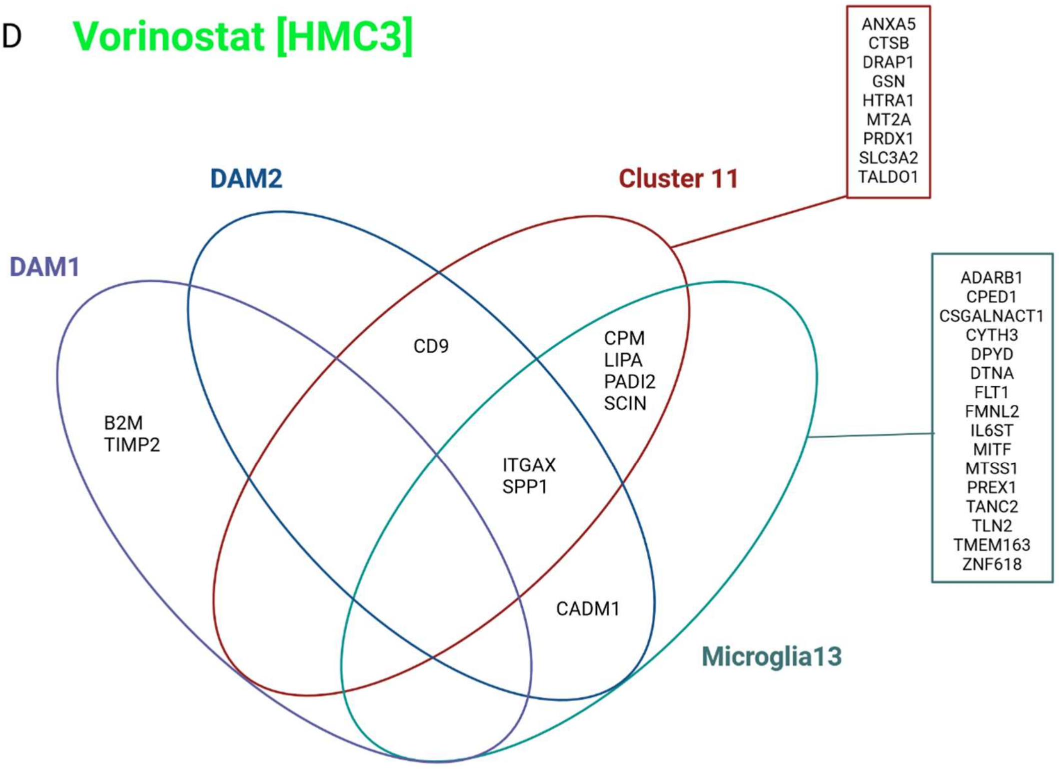

**Supplementary Figure 3. Bulk RNA-Seq of the HMC3 DAM model. A. General comparison of similarities across queried signatures of DAM-models in HMC3 microglia.** Venn diagram depicts gene markers exclusively expressed and shared across queried signatures, including DAM1 and DAM2 [3], Cluster 11 [1] and Microglia 13 [4] in Vorinostat- and Entinostat treated HMC3 microglia. Each ellipse shows specific markers for each marker set – DAM1 (purple), DAM2 (blue), Cluster 11 (red), Microglia 13 (green). Overlays of circles depict marker genes shared across different combinations of marker sets. **B. Volcano Plots depicting the distribution of differentially expressed genes from different signatures (Cluster 11 [1], Microglia 13 [4], DAM1 and DAM2 [3]) for Vorinostat and Entinostat treatment in comparison to DMSO control.** HMC3 microglia were treated for 24hrs with DMSO as control, Vorinostat (1μM) or Entinostat (10μM) followed by bulk RNA-Seq. Volcano plots depict all genes detected with all genes downregulated in blue and all genes upregulated in red, plotted based on log2FC (fold change expression) and -log10(pvalue). A selection of the top 5-10 genes upregulated for each respective marker set (Cluster 11: 89 genes, Microglia 13: 127 genes, DAM1: 10 genes, DAM2: 20 genes) is depicted. **C. Venn diagram depicting significantly induced markers across the signatures for DAM1 and DAM2 [3], Cluster 11 [1], Microglia 13 [4] in Entinostat-treated HMC3 microglia.** Each ellipse shows significantly induced markers from each marker set – DAM1 (violet), DAM2 (blue), Cluster 11 (red), Microglia 13 (green). Overlays of ellipses depict induced marker genes shared across different combinations of marker sets. **D. Venn diagram depicting significantly induced markers across the signatures for DAM1 and DAM2 [3], Cluster 11 [1], Microglia 13 [4] in Vorinostat-treated HMC3 microglia.** Each ellipse shows significantly induced markers from each marker set – DAM1 (violet), DAM2 (blue), Cluster 11 (red), Microglia 13 (green). Overlays of ellipses depict induced marker genes shared across different combinations of marker sets.

**Supplementary Figure 4**

A **General comparison across queried signatures [iMG]**

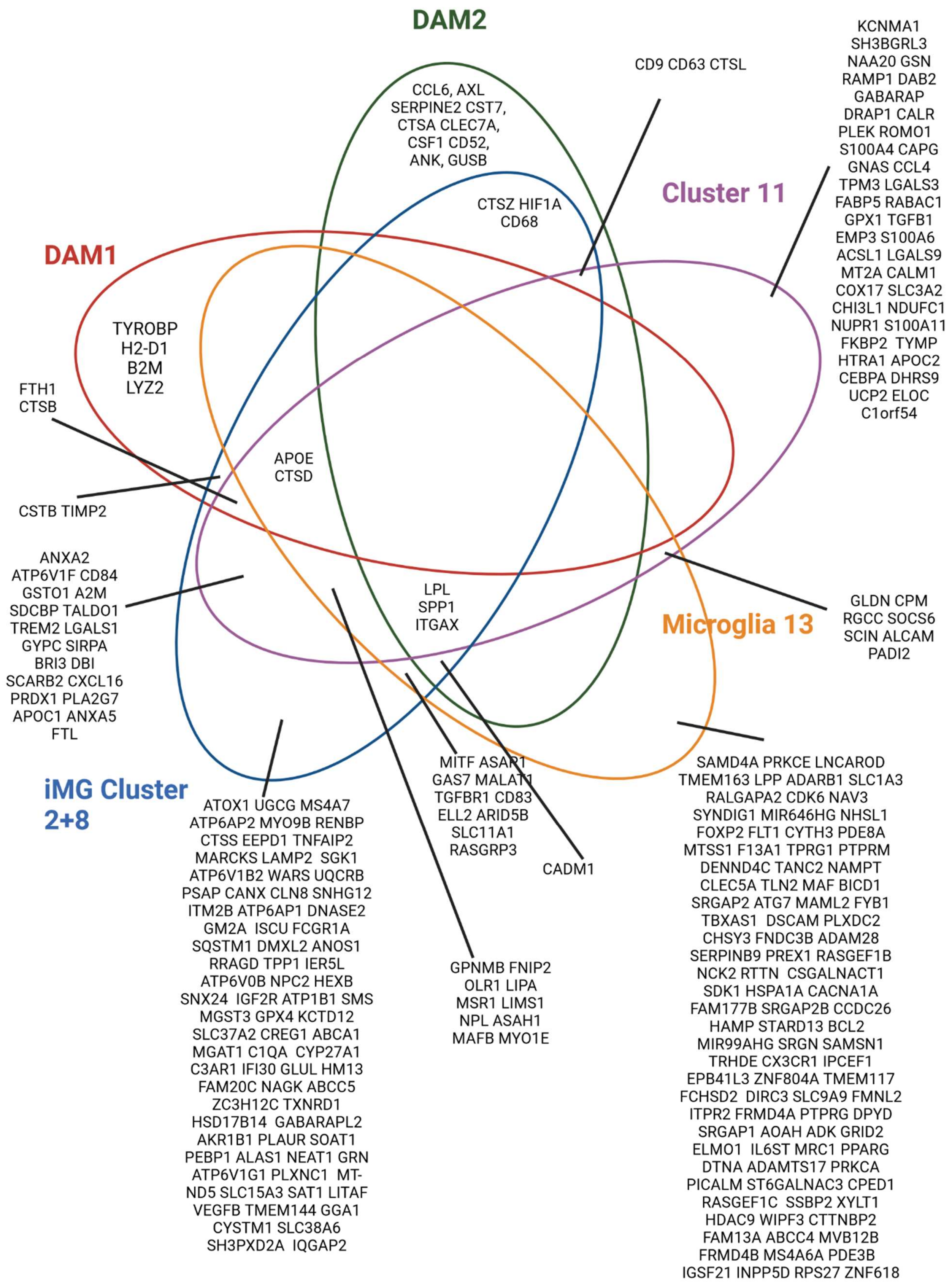

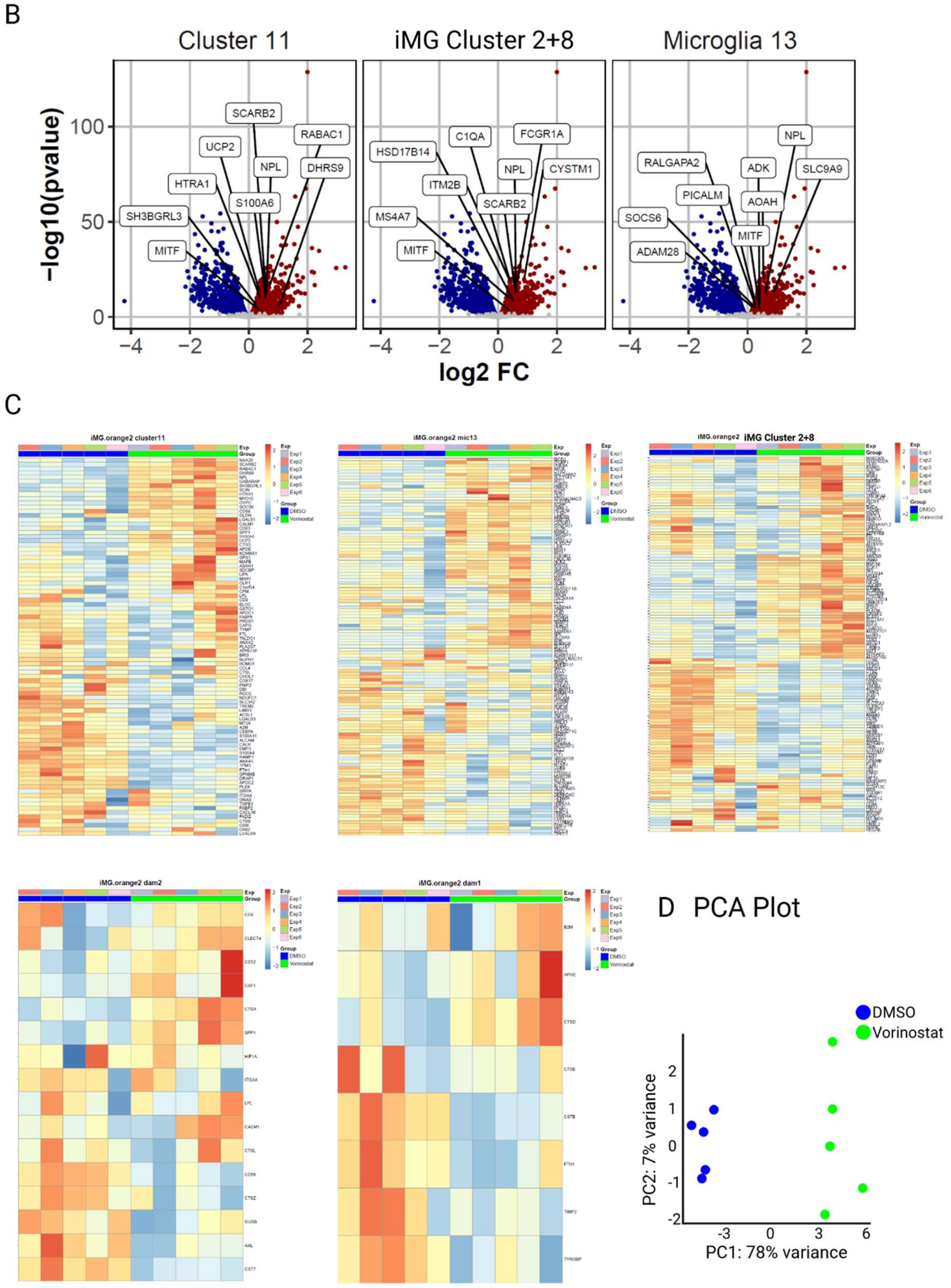
B
Cluster 11
iMG Cluster 2+8
Microglia 13
-log10(pvalue)
log2 FC
SCARB2
RABAC1
UCP2
NPL
DHRS9
HTRA1
S100A6
SH3BGRL3
MITF
C1QA
FCGR1A
HSD17B14
CYSTM1
ITM2B
MS4A7
RALGAPA2
ADK
SLC9A9
PICALM
AOAH
SOCS6
ADAM28
C
iMG.orange2 cluster11
iMG.orange2 mic13
iMG.orange2 iMG Cluster 2+8
iMG.orange2 dam2
iMG.orange2 dam1
Exp
Group
DMSO
Vorinostat
D PCA Plot
DMSO
Vorinostat
PC2: 7% variance
PC1: 78% variance

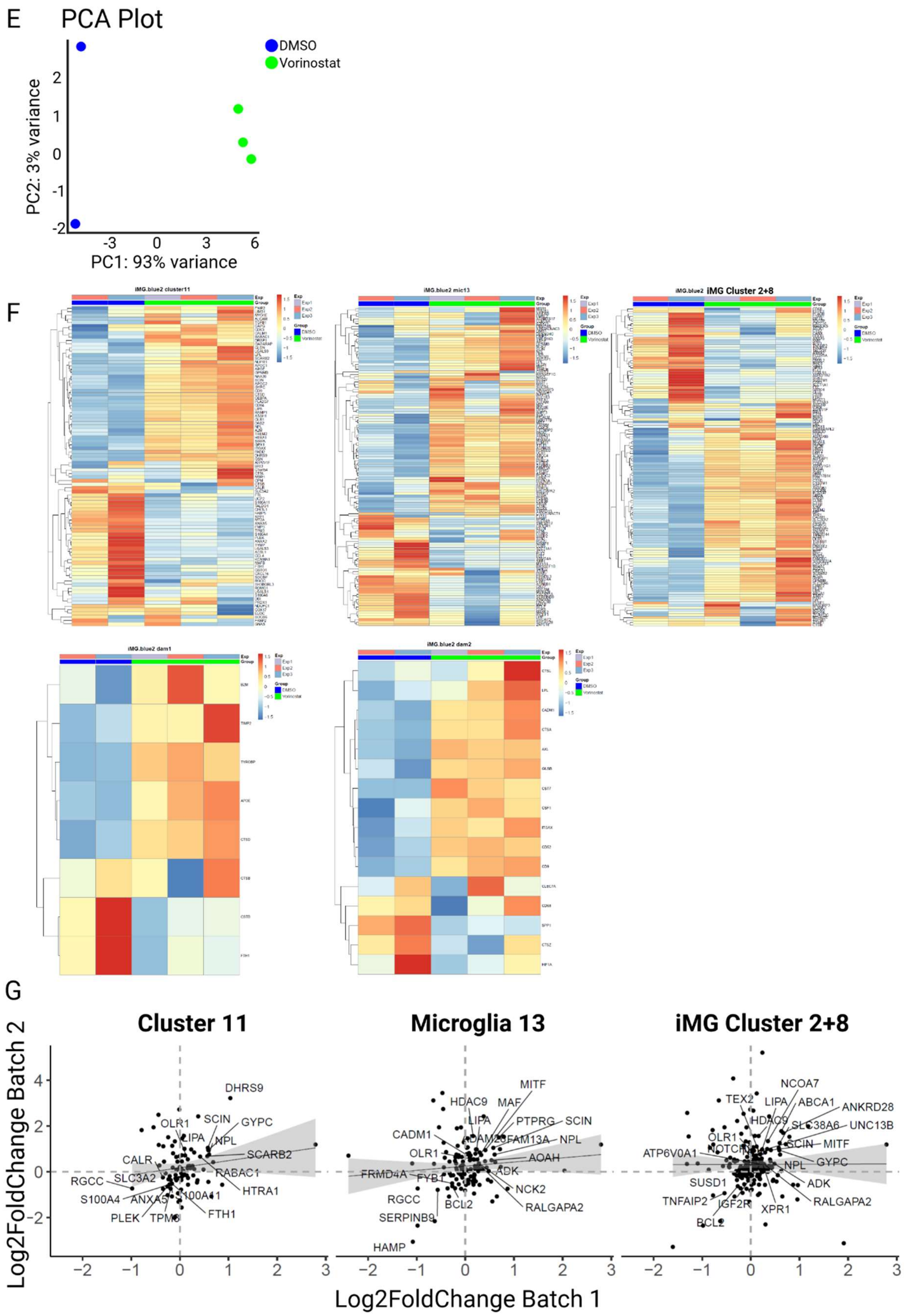

E
PCA Plot
DMSO
Vorinostat
PC2: 3% variance
PC1: 93% variance
F
iMG.blue2 cluster11
iMG.blue2 mic13
iMG.blue2 iMG Cluster 2+8
iMG.blue2 dam1
iMG.blue2 dam2
Exp
Group
Exp1
Exp2
Exp3
DMSO
Vorinostat
G
Cluster 11
Microglia 13
iMG Cluster 2+8
Log2FoldChange Batch 2
Log2FoldChange Batch 1
DHRS9
SCIN
GYPC
OLR1
LIPA
NPL
SCARB2
CALR
RABAC1
RGCC
SLC3A2
HTRA1
S100A4
ANXA5
S100A11
PLEK
TPM3
FTH1
MITF
HDAC9
MAF
PTPRG
CADM1
ADAM28
FAM13A
AOAH
FRMD4A
FYB1
ADK
NCK2
BCL2
RALGAPA2
SERPINB9
HAMP
NCOA7
TEX2
ABCA1
ANKRD28
SLC38A6
UNC13B
ATP6V0A1
SUSD1
TNFAIP2
IGF2R
XPR1

G **Signatures induced by Vorinostat across models [HMC3 & iMG]**

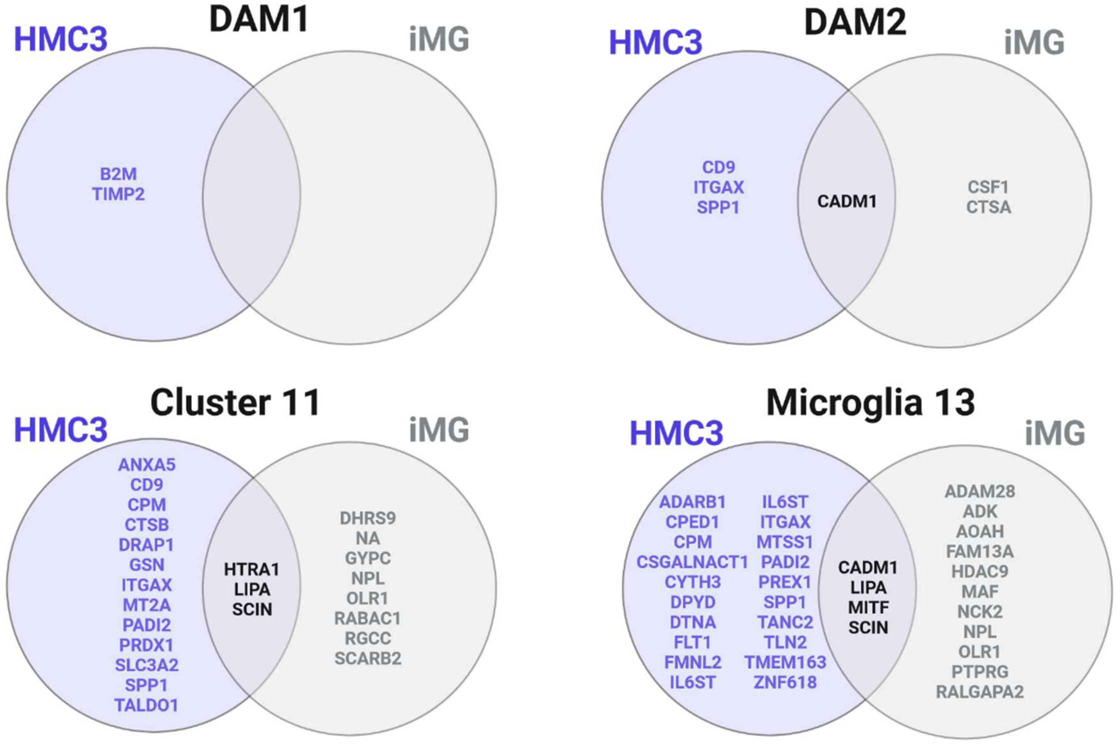


**Supplementary Figure 4. Bulk RNA-Seq of the iMG DAM model. A. General comparison of similarities across queried signatures of iPSC-derived microglia (iMG) DAM-model.** Venn diagram depicts gene markers exclusively expressed and shared across queried signatures, including DAM1 and DAM2 [3], Cluster 11 [1], Microglia 13 [4] and iMG Cluster 2+8[5] in Vorinostat- treated iMG. Each ellipse shows specific markers for each marker set – DAM1 (red), DAM2 (green), Cluster 11 (violet), Microglia 13 (yellow) and iMG Cluster 2+8 (blue). Overlays of circles depict marker genes shared across different combinations of marker sets. **B. Volcano Plots depicting the distribution of differentially expressed genes from different signatures (Cluster 11 [1], Microglia 13 [4], iMG Cluster 2+8[5]) for Vorinostat treatment in comparison to DMSO control.** iPSC-derived microglia at Day 28-29 of differentiation were treated for 24hrs with DMSO as control or Vorinostat (0.1µM) followed by bulk RNA-Seq. Volcano plots depict all genes detected with all genes downregulated in blue and all genes upregulated in red, plotted based on log2FC (fold change expression) and -log10(pvalue). A selection of the top 5-10 genes upregulated for each respective marker set (Cluster 11: 89 genes, Microglia 13: 127 genes, iMG Cluster 2+8: 134 genes) is depicted. Plots are organized from Cluster 11 (left), to Microglia 13 (middle), to iMG Cluster 2+8 (right). **C. Heatmaps showing the expression of Cluster 11** ([1]), **Microglia 13** ([4]; middle), iMG Cluster 2+8 ([5]; right), **DAM1 and DAM2** [3] **marker sets** in bulk RNA-Seq data generated 24hrs following compound treatment of iMG with DMSO (control) or Vorinostat (green; 0.1µM). Each column represents a single sample, each row a single gene represented in the respective marker set. Pairwise differential testing between DMSO control and each of the treatment conditions (Entinostat, 10µM; Vorinostat, 1µM) was conducted using a Wald test with the Benjamini-Hochberg correction (FDR alpha < 0.05). The legend represents Z scores, with lower scores indicated in red and higher scores indicated in blue. Data represents n=5 independent experiments per treatment group from one batch of iPSC-derived human microglia. **D. PCA plot of bulk RNA-Seq results from iMG treated with DMSO or Vorinostat**. Principal component analysis (PCA) was calculated on log-normalized bulk RNA-Seq data derived from compound-treated iMG following 24hrs of exposure to DMSO (control; blue) or Vorinostat (0.1µM; green). Data represents n=5 independent replicates for each of treatment group derived from one iMG batch. **E. PCA plot of bulk RNA-Seq results from a second batch of iMG treated with DMSO or Vorinostat**. Principal component analysis (PCA) was calculated on log-normalized bulk RNA-Seq data derived from compound-treated iMG following 24hrs of exposure to DMSO (control; blue) or Vorinostat (0.1µM; green). Data represents n=2 independent replicates for DMSO and n=3 replicates for Vorinostat. **F. Heatmaps showing the expression of Cluster 11** ([1]), **Microglia 13** ([4]; middle), **iMG Cluster 2+8** ([5]; right), **DAM1 and DAM2** [3] **marker sets** in bulk RNA-Seq data generated 24hrs following compound treatment of a second batch of iMG with DMSO (control) or Vorinostat (green; 0.1µM). Each column represents a single sample, each row a single gene represented in the respective marker set. Pairwise differential testing between DMSO control and each of the treatment conditions (Entinostat, 10µM; Vorinostat, 1µM) was conducted using a Wald test with the Benjamini-Hochberg correction (FDR alpha < 0.05). The legend represents Z scores, with lower scores indicated in red and higher scores indicated in blue. Data represents n=2 independent replicates for DMSO and n=3 replicates for Vorinostat. **G. Correlation analysis comparing first (orange) and second (blue) batch of iPSC-derived microglia bulk RNA-Seq datasets following treatment with DMSO or Vorinostat for 24hrs.** Data depict correlation of all genes detected by log2 fold change expression for each of the queried signatures, Cluster 11 [1], Microglia 13 [4] and iMG Cluster 2+8[5] between the two different iMG batches (orange: n=5 for each treatment; blue: n=2 for DMSO, n=3 for Vorinostat) treated with DMSO or Vorinostat (0.1µM). **H. Venn diagram depicting signatures induced by Vorinostat across both DAM models in HMC3 microglia and iPSC-derived human microglia (iMG).** Each venn diagram depicts the genes induced in either HMC3 (left,blue) or iMG (right, grey) or in both cell models (overlapping center part) for DAM1 (upper left) or DAM2[3] (upper right), Cluster 11 [1](lower left) or Microglia 13[4] (lower right).

### Supplementary Figure 5

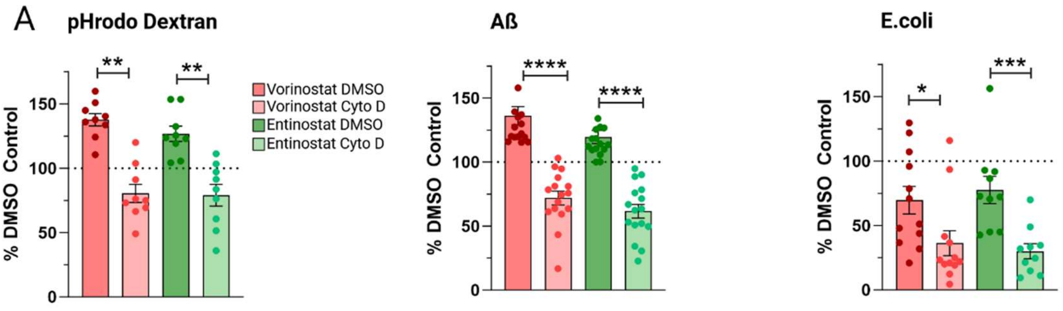


**Supplementary Figure 5. A. Complete figures of macropinocytosis and phagocytosis assays following compound-treatment (Vorinostat, Entinostat, DMSO as control) using different substrates (pHrodo Dextran, Aß, pHrodo E.coli) including negative controls (Cytochalasin D treated cells).** For all three assays, HCM3 microglia were pretreated with respective compounds (Vorinostat, Entinostat) or DMSO as control for 24hrs, subsequently exposed to pHrodo-labeled Dextran or Aß1-42 or pHrodo E.coli either containing 5µM Cytochalasin D (Sigma-Aldrich, Cat#: C8273) as a negative control for phagocytosis, or DMSO as control for Cytochalasin D treatment. After 1hr the uptake of the respective substrate was assessed using flow cytometry. Individual experiments are depicted as individual dots in the bar graphs depicting mean ± SEM (Camptothecin – orange; Narciclasien – blue; Torin2 - purple). Phagocytosis was normalized to % DMSO control and for statistical analysis, log-fold change values of compound- and Cytochalasin D -treated cells in comparison to compound-treated samples was analyzed using Mann-Whitney U test. *p.adj ≤ 0.05; **p.adj ≤ 0.01; ***p.adj ≤ 0.001; ****p.adj ≤ 0.0001.
